# Impaired LAIR-1-mediated immune control due to collagen degradation in fibrosis

**DOI:** 10.1101/2021.12.07.471103

**Authors:** Tiago Carvalheiro, Wioleta Marut, M. Inês Pascoal Ramos, Samuel García, Devan Fleury, Alsya J. Affandi, Aniek S. Meijers, Barbara Giovannone, Ralph G. Tieland, Eline Elshof, Andrea Ottria, Marta Cossu, Matthew L. Meizlish, Tineke Veenendaal, Meera Ramanujam, Miguel E. Moreno-García, Judith Klumperman, Nalan Liv, Timothy R.D.J. Radstake, Linde Meyaard

## Abstract

Tissue repair is disturbed in fibrotic diseases like systemic sclerosis (SSc), where the deposition of large amounts of extracellular matrix components such as collagen interferes with organ function. LAIR-1 is an inhibitory collagen receptor highly expressed on tissue immune cells. We questioned whether in SSc, impaired LAIR-1-collagen interaction is contributing to the ongoing inflammation and fibrosis.

We found that SSc patients do not have an intrinsic defect in LAIR-1 expression or function. Instead, fibroblasts from SSc patients deposit disorganized collagen products *in vitro*, which are dysfunctional LAIR-1 ligands. This can be mimicked in healthy fibroblasts stimulated by soluble factors that drive inflammation and fibrosis in SSc and is dependent of matrix metalloproteinases and platelet-derived growth factor receptor signaling.

In support of a non-redundant role of LAIR-1 in the control of fibrosis, we found that LAIR-1-deficient mice have increased skin fibrosis in response to repeated injury and in the bleomycin mouse model for SSc. Thus, LAIR-1 represents an essential control mechanism for tissue repair. In fibrotic disease, excessive collagen degradation may lead to a disturbed feedback loop. The presence of functional LAIR-1 in patients provides a therapeutic opportunity to reactivate this intrinsic negative feedback mechanism in fibrotic diseases.

## Introduction

The extracellular matrix (ECM) is a dynamic structure, in which production and deposition are finely balanced with degradation and removal, regulated through pro-fibrotic factors, matrix metalloproteinases (MMPs) and other collagenases. Tightly controlled ECM homeostasis is essential for healthy connective tissues^1,2^. Tissue repair after injury entails a cascade of inflammation, repair and ultimately resolution. Tissue resident macrophages can sense ECM composition. Signals from damaged tissue initiate a dynamic process that involves the recruitment of immune cells, which help to induce fibroblast differentiation into myofibroblasts, which in turn deposit ECM to repair the tissue. Finally, myofibroblasts are eliminated and the ECM is degraded and remodeled, in order to resolve the repair process^2,3^. Fibrosis is an abnormal reparative process that is characterized by excessive ECM deposition without reciprocal degradation^4^. This progressive accumulation of ECM results in remodeling of the tissue architecture and can lead to organ failure^5,6^. The fibrotic scar consists predominantly of fibrillar collagens (types I and III), fibronectin and basement-membrane proteins such as type IV collagen and laminin^7–9^. Complex cellular and molecular processes drive fibrosis, with transforming growth factor β (TGFβ) family proteins playing a central role by stimulating the synthesis of ECM proteins by effector cells such as myofibroblasts^10,11^.

Systemic sclerosis (SSc) is a prototypical immune-mediated fibrotic disease, characterized by vasculopathy, inflammation and fibrosis of the skin and internal organs. The etiology of SSc is largely unknown and patients can present with a wide range of clinical manifestations, resulting in high morbidity and mortality^12^. Typically, immune cell infiltration into skin and internal organs occurs early in the disease, prior to the onset of fibrosis. Immune mediators such as TGFβ - namely TGFβ2, type I interferon (IFN), platelet-derived growth factor (PDGF), IL-6 and type 2 cytokines (including IL-4, IL-10 and IL-13) are considered central to the pathogenesis of SSc^13,14^. In addition, the chemokine CXCL4 is increased in the circulation and skin of SSc patients and correlates with the presence and progression of complications such as lung fibrosis and pulmonary arterial hypertension^15^.

ECM proteins interact with different cell types through cell surface receptors, including integrins, receptor tyrosine kinases, and immunoglobulin-like receptors^16^. Leukocyte associated immunoglobulin-like receptor-1 (LAIR-1; CD305) is an Immunoreceptor Tyrosine-based Inhibitory Motif (ITIM)-bearing inhibitory receptor for collagen, broadly expressed on most immune cells^17,18^. Activation of LAIR-1 *in vitro* potently inhibits diverse immune cells such as T cells, NK cells^17,19–21^, B cells^22^ plasmacytoid dendritic cells (pDC)^23,24^ and monocytes^25–27^. LAIR-1 inhibitory function can be regulated at the membrane expression level and by the natural agonists soluble-LAIR-1 (sLAIR-1), a shed form of LAIR-1, and LAIR-2. LAIR-2 is a soluble protein, homologous to human LAIR-1, lacking the transmembrane and intracellular domain, with a higher affinity for collagens than LAIR-1, thereby blocking LAIR-1 ligation to collagen^28^.

LAIR-1 is expressed on immune cells in healthy skin, where it can interact with collagens^18,26^. It recognizes multiple collagens and collagen-domain containing proteins. Collagens have multiple functional binding sites for LAIR-1, the glycine-proline-hydroxyproline content of collagen domains appears to be the dominant determinant for LAIR-1 binding^18,29^. In fibrotic tissues such as those in SSc patients, there should be abundant amounts of collagen to be sensed by LAIR-1 on tissue immune cells, yet inflammation continues. We here questioned whether in SSc, impaired LAIR-1-collagen interaction is contributing to the ongoing inflammation and fibrosis.

## Results

### LAIR-1 expression and its intrinsic function is normal in SSc patients

We first investigated whether the function of LAIR-1 is impaired in patients with SSc. We determined LAIR-1 expression on circulating immune cells from SSc patients and healthy controls (HC) by flow cytometry. LAIR-1 expression at the cell surface was comparable between SSc patients and HC on type-2 conventional dendritic cells (cDC2), plasmacytoid dendritic cells (pDCs), CD4^+^ and CD8^+^ T cells, NK cells and B cells and on most monocyte subsets. In classical monocytes, LAIR-1 expression was significantly increased in SSc patients compared to HC (Figure 1A). As described before^26^, LAIR-1 was not expressed on type-1 conventional dendritic cells (cDC1) (data not shown).

**Figure 1.**
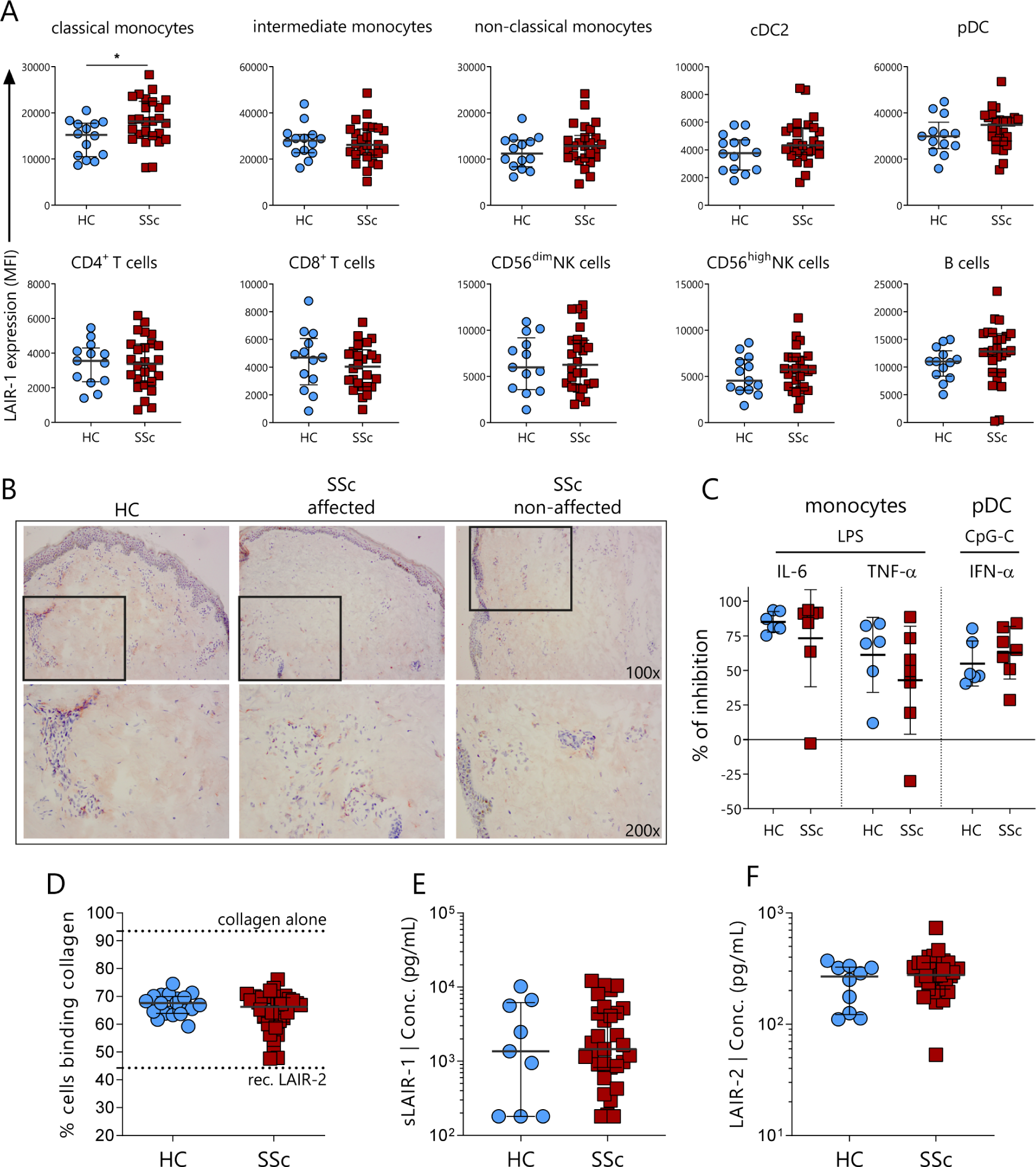
LAIR-1 is expressed on circulating peripheral blood cells and in the skin of SSc patients and its intrinsic function is normal. **(A)** LAIR-1 expression on circulating immune cells (classical, intermediate and non-classical monocytes, cDC2, pDC, CD4^+^ and CD8^+^ T cells, CD56^bright^ and CD56^dim^ NK cells and B cells) from HC (n=14) and SSc patients (n=26) determined using flow cytometry. **(B)** Representative images of LAIR-1 expression (shown as red precipitate) in affected and non-affected skin from HC skin (n=3) and SSc patients (n=3), assessed by immunohistochemistry. **(C)** LAIR-1 function in monocytes and pDC from HC (n=6) and SSc patients (n=7). Isolated cells were pre-treated with a plate bound LAIR-1 agonist mAb (clone Dx26) or isotype control antibody. Monocytes were subsequently stimulated (24 hours) with LPS – TLR4 ligand, while pDCs were stimulated (24 hours) with CpC-C – TLR9 ligand. Production of IL-6 and TNF-α by monocytes and IFN-α by pDCs was determined by ELISA. Percentage (%) inhibition of cytokine production by LAIR-1 mAb is shown. **(D)** Effect of serum from HC and SSc patients to block LAIR-1 – collagen interaction. FITC labelled collagen was incubated either with sera from HC (n=20) or SSc patients (n=36) and the binding capacity of collagen to bind K562 cells overexpressing LAIR-1 was determined by flow cytometry. Collagen alone and blocking with recombinant LAIR-2 (40 µg/mL) are shown as controls with the dashed lines. Serum levels of **(E)** soluble LAIR-1 and **(F)** LAIR-2 in circulation of HC (n=10) and SSc patients (n=30) determined using ELISA. Results are represented as mean ± SD. Differences were considered statistically significant when *p < 0.05 (Mann–Whitney test).

In affected skin of SSc patients, available profiling data by array [GSE95065] show that *LAIR1* gene expression was increased in the skin of SSc patients (Supplemental Figure 1A), probably reflecting an increase in immune cells, given the strong correlation between *LAIR1* expression and *PTPRC* (gene encoding CD45) and moderate correlation with the macrophage marker *CD68* (Supplemental Figure 1B). By immunohistochemistry, we confirmed that LAIR-1 was expressed in skin from HC and SSc patients, both in affected and non-affected skin (Figure 1B). Thus, LAIR-1 is expressed on circulating immune cells and skin of patients with SSc.

We next investigated whether LAIR-1 on immune cells from SSc patients retains its capacity to inhibit cellular activation. Monocytes and pDC from HC and SSc patients were stimulated with TLR ligands together with an antibody that acts as a LAIR-1 agonist. LAIR-1 activation in both groups caused a similar inhibition of IL-6 and TNF-α production by monocytes and IFN-α production by pDC (Figure 1C). Thus, the intrinsic inhibitory function of LAIR-1 is preserved in SSc patients.

We further explored whether soluble mediators in the circulation of SSc patients could block collagen interaction with cellular LAIR-1. For this purpose, labelled collagen was incubated with serum from SSc patients or HC after which collagen binding to a LAIR-1 overexpressing line was determined by flow cytometry. Serum from SSc patients and HC blocked collagen binding to LAIR-1 to a similar degree (Figure 1D, Supplemental Figure 2A). Consistent with this finding, circulating concentrations of sLAIR-1 (Figure 1E) and LAIR-2 (Figure 1F) were not different between SSc patients and HC. Altogether, these experiments indicate that LAIR-1 is normally expressed, has preserved inhibitory capacity, and does not encounter higher levels of known LAIR-1 antagonists in SSc patients compared to HC.

### LAIR-1 binds to *in vitro* and *in vivo* fibrotic ECM

We have previously shown that LAIR-1 is able to bind to collagen in healthy skin^18^. The fibrotic collagen present in the skin of SSc patients is structurally different from that found in healthy skin, with dense, aligned dermal collagen bundles^30^, which potentially could affect LAIR-1 binding. However, we observed that LAIR-1 binding to both HC and SSc skin samples was comparable (Figure 2A).

**Figure 2.**
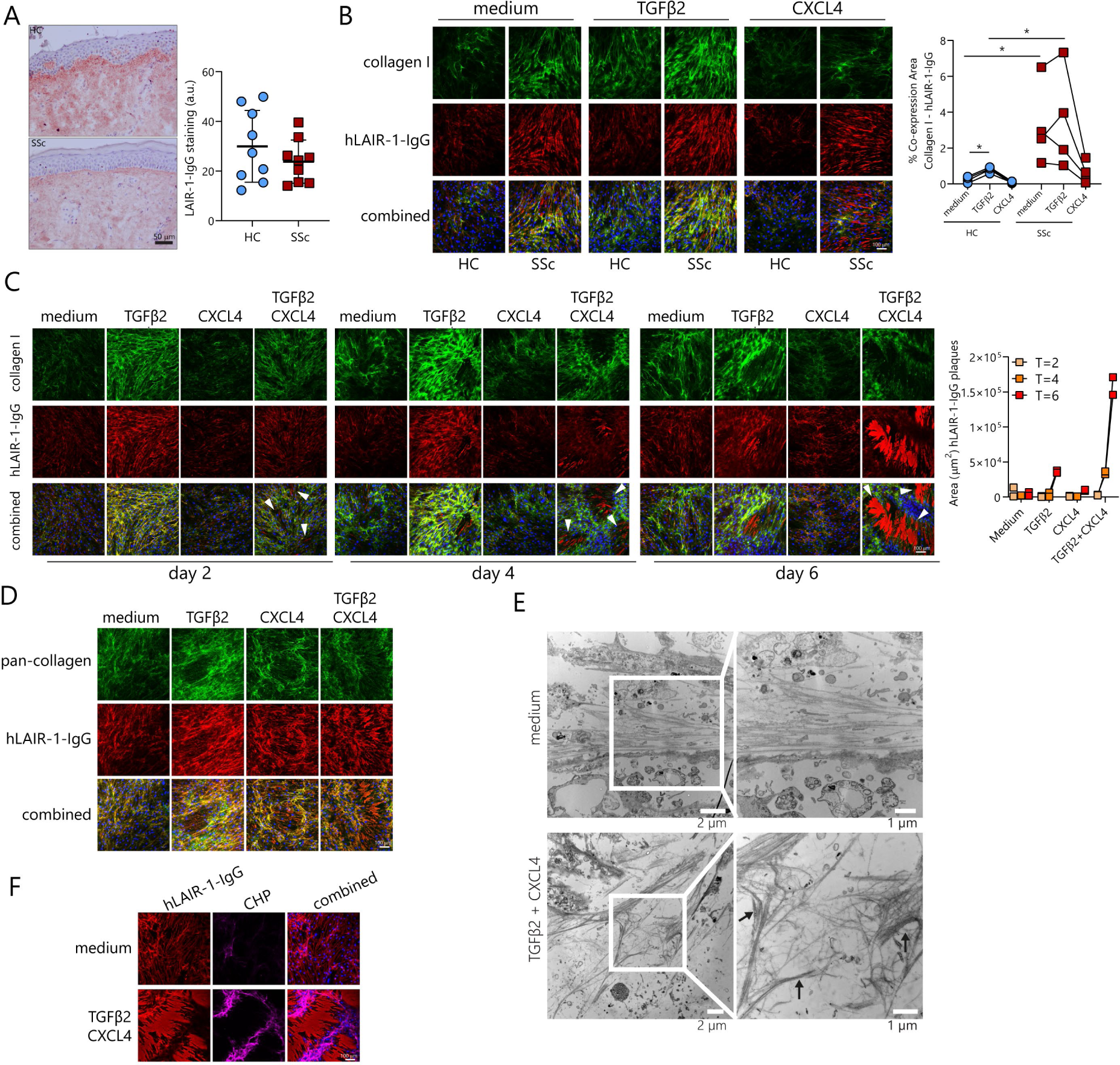
LAIR 1 binds to *in vitro* and *in vivo* fibrotic ECM. **(A)** Representative immunohistochemistry staining and quantification of the hLAIR-1-IgG fusion protein (shown in red) and counterstained with hematoxylin in the skin of HC (n=9) and SSc patients (n=9). **(B)** Representative images of immunofluorescence (IF) staining with collagen I antibody (green), hLAIR-1-IgG fusion protein (red) and DAPI (blue) of ECM produced by dermal fibroblasts from HC (n=4) and SSc patients (n=4) stimulated with either medium, TGF-β2 or CXCL4. Quantification of % area with co-expression of Collagen I and hLAIR-1-IgG is shown. **(C)** Representative IF staining with collagen I antibody (green), hLAIR-1-IgG fusion protein (red) and DAPI (blue) of ECM produced by HC dermal fibroblasts stimulated with medium, TGF-β2, CXCL4 or TGF-β2 combined with CXCL4. White arrows indicate deposits of LAIR-1 binding structures (representative of n=2). The area (µm^2^) of hLAIR-1-IgG plaques is quantified **(D)** Representative IF staining with hLAIR-1-gG fusion protein (red), pan collagen antibodies (mix of collagen I, II, III, IV and V antibodies) (green) and DAPI (blue) of ECM produced by HC dermal fibroblasts stimulated with medium, TGF-β2, CXCL4 or TGF-β2 combined with CXCL4. (representative of n=3) **(E)** Transmission electron microscopy (TEM) images showing the high-resolution structure and organization of collagen fibers in the ECM produced by HC in medium control (upper panel) or after stimulation with TGFβ2 plus CXCL4 (lower panel). Black arrows depict dense bundles of collagen fibers observed in TGFβ2 plus CXCL4 treated fibroblasts. Scale bars are 2 µm and 1 µm for the left and the right images in both conditions, respectively. **(F)** Representative IF staining with collagen hybridizing peptide (CHP) (purple) for denatured collagen, hLAIR-1-IgG fusion protein (red) and DAPI (blue) on ECM produced by HC in medium condition or after stimulation with TGFβ2 combined with CXCL4 (n=4). Representative IF images were acquired on a confocal microscope with a 20x magnification. Results are represented as mean with SD. Differences were considered statistically significant when *p < 0.05 (Mann–Whitney test HC vs SSc patients groups, paired samples tested with RM one-way ANOVA, followed by Fisher’s LSD). Results are represented as mean with SD.

Fibroblasts in the skin of SSc patients are constantly exposed to inflammatory and pro-fibrotic factors, such as TGFβ and CXCL4^13^. We mimicked these circumstances *in vitro* by culturing dermal fibroblasts obtained from HC and SSc patients in the presence of TGFβ or CXCL4 and visualized collagen I deposition by immunofluorescence. TGFβ treatment induced enhanced collagen I production compared to untreated (medium) cultures, and collagen was deposited in characteristic aligned fibers, both in cultures of fibroblasts from SSc patients and from HC. In contrast, CXCL4 stimulation resulted in more disorganized collagen I, particularly in fibroblasts from SSc patients (Figure 2B). We assessed LAIR-1 binding to the deposited ECM using a hLAIR-1-IgG fusion protein^18^. We observed that LAIR-1 efficiently bound to the ECM produced by fibroblasts from both HC and SSc patients, both in non-stimulated and stimulated conditions. However, when compared to HC, SSc dermal fibroblasts stimulated with CXCL4 produced an increased amount of dense, non-fibrillar shaped structures that bound LAIR-1 but did not bind collagen I antibodies (Figure 2B). Thus, SSc fibroblasts stimulated with CXCL4, produce an ECM structure that is not recognized by collagen I antibodies and binds LAIR-1.

We next determined whether HC dermal fibroblasts stimulated with the potent pro-fibrotic combination of TGFβ and CXCL4 induced similar LAIR-1 binding structures as observed in SSc fibroblasts treated with CXCL4 only. Indeed, after stimulation with TGFβ and CXCL4, HC fibroblasts deposited similar dense structures binding LAIR-1 but not collagen I antibodies (Figure 2C). The deposition of these structures increased evidently over time and was preferentially localized to decellularized areas (Figure 2C). Co-staining of the ECM with a pan-collagen mix of antibodies (recognizing collagen I, II, III, IV and V) revealed these structures as collagen, even though they did not have a typical fibrillar collagen conformation (Figure 2D). In addition, we confirmed that these structures do not bind to human Signal inhibitory receptor on leukocytes-1 (hSIRL-1), a protein with homology to LAIR-1 (Supplemental Figure 3A).

To study these fibrotic collagen structures at the nanoscale, fibroblasts were stimulated with TGFβ2 and CXCL4 for transmission electron microscopy (TEM). High-resolution TEM images showed that the ECM of the non-stimulated (medium) condition was composed mostly of aligned collagen fibers that were homogenously interspaced and run parallel to each other (Figure 2E). In contrast, collagen fibers in the ECM produced by fibroblasts stimulated with TGFβ2 and CXCL4 were more frequently disorganized, had unidirectional orientation, and sometimes formed dense bundles (Figure 2E-black arrows and Supplemental Figure 4A-B). The disrupted organization of collagen fibers upon TGFβ2 and CXCL4 treatment can possibly be a consequence of the degradation of collagen fibers or collagen associated proteins responsible for their organization.

Since degraded and fragmented collagen triple helices may become unstable and spontaneously unfold, leaving denatured collagen fragments, we next assessed whether the LAIR-1 binding structures correspond to denatured collagen. We used a collagen hybridizing peptide (CHP), which specifically binds to denatured collagen strands^31^. CHP did bind in cell-dense areas where collagen is actively produced but did not co-localize with the LAIR-1-binding collagen structures (Figure 2F). This suggests that the collagen triple helical conformation is intact in the degraded fibers, in agreement for a need of a triple helical conformation for LAIR-1 binding^18,29^. Thus, fibroblasts stimulated with TGFβ and CXCL4 deposit dense structures consisting of degraded triple helical collagen in decellularized areas that specifically bind LAIR-1.

### PDGF receptor signaling and Matrix Metalloproteinases are required for deposition of fibrotic LAIR-1 binding structures

The areas where the LAIR-1-binding dense collagen structures were deposited in *in vitro* cultures were devoid of cells (Figure 2C). By live cell imaging, we observed that the combination of TGFβ with CXCL4 drastically altered the motility of both HC and SSc dermal fibroblasts. Instead of forming an intact cellular monolayer, fibroblasts became highly motile, losing contact inhibition and forming multicellular layers, while leaving behind large decellularized areas (Figure 3A and Supplemental video 1-2).

**Figure 3.**
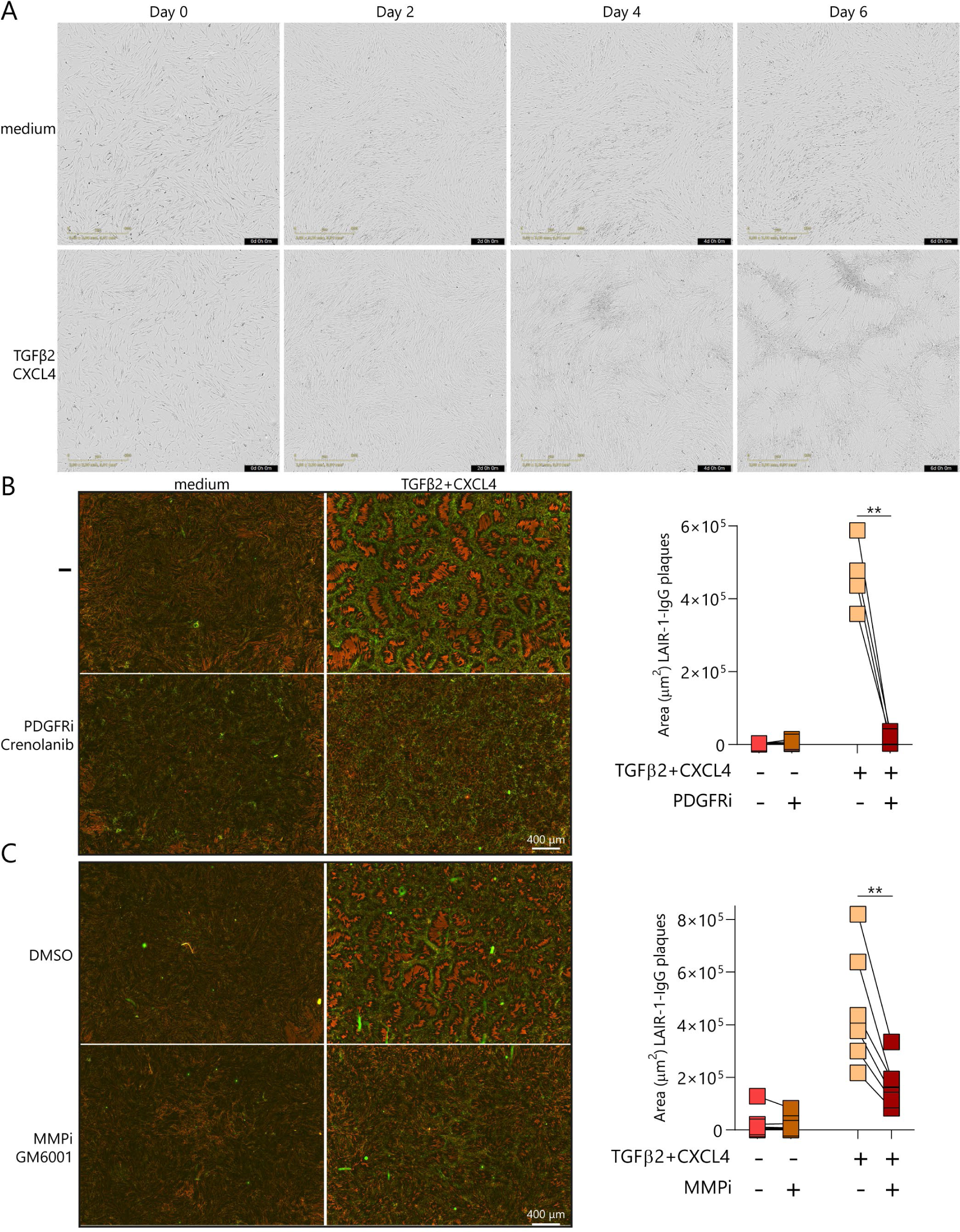
PDGF receptor signaling and Matrix Metalloproteinases are required for deposition of fibrotic LAIR-1 binding structures. **(A)** HC fibroblasts were stimulated with TGF-β2 combined with CXCL4 and imaged during 6 days on a Incucyte live system. Representative phase contrast images (4x magnification) are shown (n=3). HC fibroblasts stimulated with TGF-β2 in combination with CXCL4 **(B)** in the presence or absence of 1 µM PDGF receptor inhibitor (Crenolanib) (n=4) or **(C)** in the presence of 100 µM MMP inhibitor (GM-6001) or DMSO as negative control (n=6). Samples were stained for immunofluorescence with hLAIR-1-gG fusion protein (red) and pan collagen antibodies (green). Immunofluorescence images were acquired on Incucyte using a 10x magnification and representative tiles of 16 images 4×4 are shown. The area (µm^2^) of hLAIR-1-IgG plaques is quantified. Statistically significant differences were considered when **p < 0.01 (paired-t test).

Since PDGF signaling regulates proliferation and cell motility ^32^, we next assessed whether this pathway could be involved in the production of the fibrotic LAIR-1 binding structures. Indeed, PDGF receptor inhibition in fibroblasts stimulated with TGFβ plus CXCL4 reduced cell motility and prevented the formation of multicellular layers of fibroblasts and the occurrence of decellularized areas (Supplemental video 3-6), as well as of the deposition of LAIR-1-binding structures (Figure 3B). Thus, PDGF-mediated fibroblast activation is required for the formation of these fibrotic collagen structures.

Next, we tested whether matrix metalloproteinases (MMP) were involved in the generation of the fibrotic LAIR-1-binding structures. Inhibition of MMP activity in HC dermal fibroblasts stimulated with TGFβ plus CXCL4 reduced the amount of LAIR-1-binding ECM structures (Figure 3C), and decreased cell motility (Supplemental video 7-10). Thus, the deposition of the LAIR-1-binding fibrotic structures requires cell motility and loss of cell contact inhibition mediated by PDGF receptor signaling and collagen degradation by MMPs.

### Fibrotic collagen products have impaired capacity to activate LAIR-1

While the fibrotic collagen structures that we identified bind to hLAIR-1-IgG, we wondered whether they are functional ligands that can activate cellular LAIR-1. To investigate the effect of degraded collagen on LAIR-1 activation, we first used gelatin, which is a mixture of peptides and proteins produced by partial hydrolysis of collagen, resulting in small collagen fragments. hLAIR-1-IgG bound in a dose dependent manner to plate-bound purified collagen I and, to a lesser extent to gelatin (Figure 4A). Next, to assess the capacity of gelatin to activate LAIR-1, we used a NFAT-GFP reporter cell line transfected with LAIR-1-CD3ζ, which expresses Green Fluorescent Protein (GFP) when LAIR-1 is activated^18^. Optimal LAIR-1-CD3ζ-induced GFP production was achieved at ∼6 µg/mL of immobilized collagen I, while lower concentrations or higher concentrations resulted in a lower percentage of GFP expressing cells (Figure 4B). In contrast, gelatin did not induce GFP production at any of the concentrations tested, even at concentrations where hLAIR-1-hIg effectively bound (Figure 4A-B). These results suggest that collagen degraded by partial hydrolysis can bind LAIR-1 but cannot activate LAIR-1.

**Figure 4.**
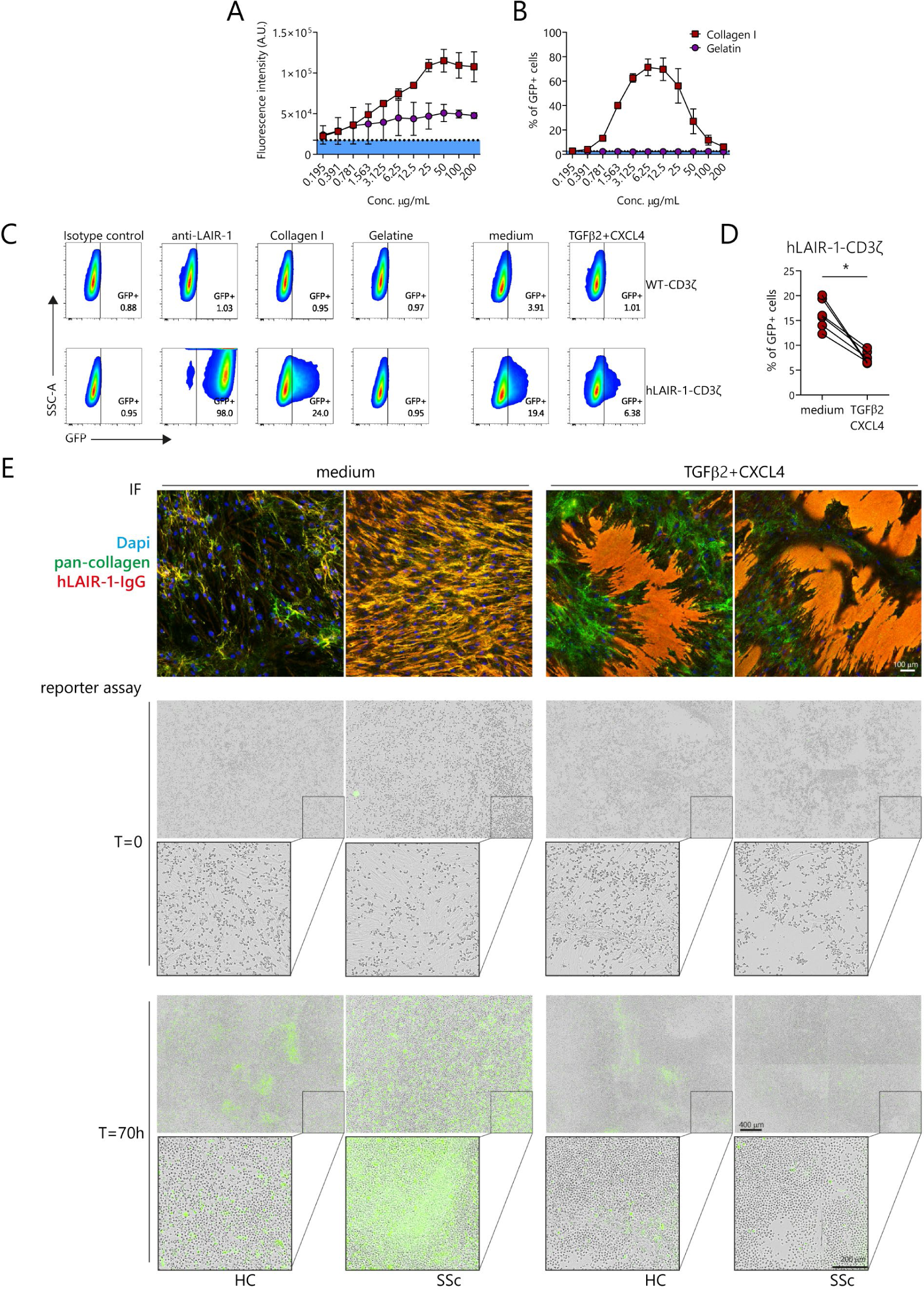
Fibrotic disorganized collagen products have impaired capacity to activate LAIR-1. **(A)** Fluorescent hLAIR 1-IgG binding and **(B)** LAIR-1 response using a NFAT GFP reporter cell line transfected with hLAIR-1 and CD3ζ chimera (hLAIR-1-CD3ζ) to plate bound purified human Collagen I, gelatin from bovine skin and BSA (bovine serum albumin) as unspecific control (shown in blue shade). **(C)** Response (% of GFP+ cells measured by flow cytometry) of hLAIR-1-CD3ζ or control WT-CD3ζ parental cells to a mouse IgG1 isotype control (negative control), anti-LAIR-1 agonistic mAb (clone Dx26) (positive control), human purified collagen I (5 µg/mL), gelatin from bovine skin (5 µg/mL) and **(D)** to ECM produced by dermal HC dermal fibroblasts unstimulated or upon stimulation with TGFβ2 plus CXCL4 during 13 days (n=6). **(E)** hLAIR1-CD3ζ reporter cells were incubated on ECM produced by SSc patients (n=4) and HC (n=4) dermal fibroblasts unstimulated or upon stimulation (6 days) with TGFβ2 combined with CXCL4 and GFP expression (represented in green) was evaluated on the Incucyte imaging system during 70 hours. Paired samples were used to immunofluorescence staining for hLAIR-1-gG fusion protein (red), pan collagen antibodies (green) and DAPI (blue) to confirm the deposit of LAIR-1-binding ECM structures. Differences were considered statistically significant when *p < 0.05 (paired-t test).

To investigate whether the fibrotic ECM structures that we identified are functional ligands for LAIR-1, we performed the same reporter cell assay on ECM produced by fibroblasts stimulated with TGFβ and CXCL4. We were able to trigger GFP expression in approximately 24% of LAIR-1-CD3ζ reporter cells using purified human collagen immobilized in the same culture vessel used in our ECM culture system (Figure 4C). ECM produced by unstimulated fibroblasts triggered GFP expression in 16.25±3.07% of the LAIR-1-CD3ζ reporter cells. However, ECM produced by TGFβ plus CXCL4 stimulated cells only resulted in 7.54±1.27% of GFP+ LAIR-1-CD3ζ reporter cells, which is a 54% reduction in the capacity to activate LAIR-1 (Figure 4C-D). We additionally performed a live reporter cell assay on ECM produced by fibroblasts from HC and SSc patients after stimulation with TGFβ plus CXCL4. After 70 hours of culture, GFP positive reporter cells were detected in areas where intact collagen structures were present. In contrast, in the decellularized regions, where degraded hLAIR-1-IgG binding collagen structures are deposited, LAIR-1-CD3ζ reporter cells were not induced to express GFP (Figure 4E and Supplemental video 11-14). This indicates that, despite having the capacity to bind LAIR-1, these degraded collagen structures did not activate LAIR-1. The production of degraded collagen in fibrotic conditions such as SSc could thus lead to incapacitated LAIR-1 inhibitory function and immune cell control.

### Absence of LAIR-1 leads to increased bleomycin-induced skin fibrosis in the SSc mouse model

To investigate if the absence of functional LAIR-1 signaling would indeed lead to enhanced fibrosis, we subjected *Lair1^-/-^* mice to the bleomycin-induced fibrosis model. This is a well-established mouse model of SSc, in which daily subcutaneous injections of bleomycin are used to induce skin fibrosis^33^. The model course can be divided into an inflammatory phase (7-15 days of bleomycin injections) and a fibrotic phase (2-4 weeks of bleomycin injections) (Figure 5A).

**Figure 5.**
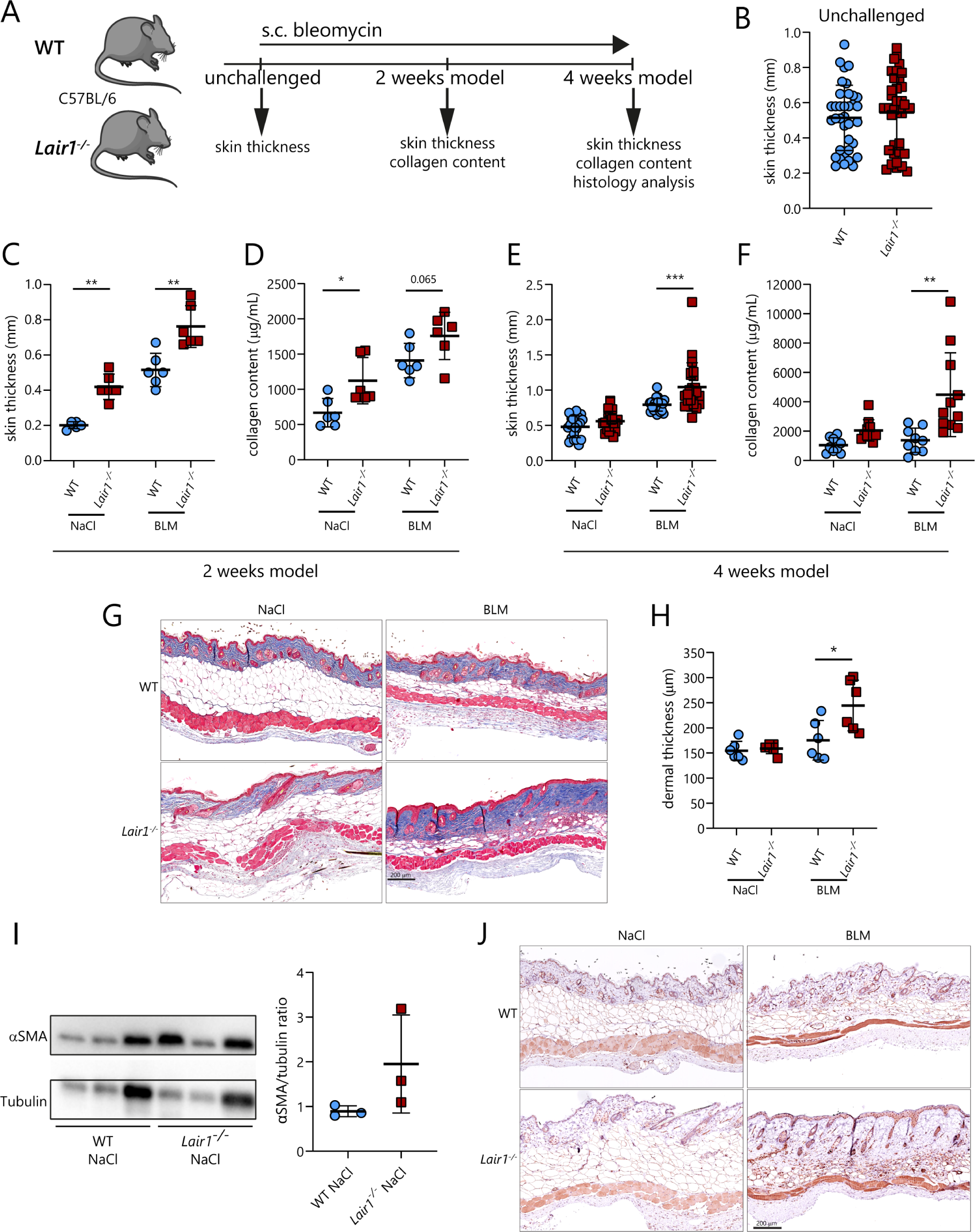
Absence of LAIR-1 leads to increased bleomycin-induced skin fibrosis in the SSc mouse model. **(A)** WT or *Lair1^-/-^* mice were subcutaneously injected with bleomycin (BLM) or saline (NaCl) control for 2 weeks or 4 weeks. **(B)** Skin thickness was measured by caliper in unchallenged WT or *Lair1^-/-^* mice (36 mice per group, 3 independent experiments). **(C)** Caliper skin thickness, **(D)** αSMA protein expression by western blot and **(E)** collagen content was determined after 2 weeks of treatment (6 mice per group, 3 per group for western blot, 1 independent experiment). In the 4 weeks treatment model **(F)** caliper skin thickness (21-23 mice per group, 4 independent experiments) and **(G)** collagen content was measured (9-11 mice per group, 2 independent experiments). From the affected skin, **(H)** representative images of Masson’s trichrome staining of affected skin and quantification (n=6 per group, 1 independent experiment) are shown, as well as **(I)** representative histological αSMA staining (n=6 per group, 1 independent experiment). Data are shown as mean ± SD. Statistically significant differences were considered when *p < 0.05, **p < 0.01, ***p < 0.001 (Mann– Whitney test WT vs *Lair1^-/-^* groups).

Untreated *Lair1^-/-^* mice did not display signs of skin fibrosis, as no differences were observed in skin thickness between *Lair1^-/-^*and WT mice measured by caliper (Figure 5B), and by histology (Supplemental Figure 5A-B). To determine whether LAIR-1 regulates the fibrotic process triggered upon bleomycin treatment, we subcutaneously injected bleomycin or sodium chloride (NaCl) as control in *Lair1^-/-^* or WT mice (Figure 5A). After 2 weeks of bleomycin treatment (inflammatory phase) *Lair1^-/-^* mice exhibited increased skin thickness compared to WT mice. Interestingly, *Lair1^-/-^* mice also showed increased skin thickness and higher αSMA protein expression compared to WT mice after repeated injection of the vehicle NaCl (Figure 5C-D), suggesting that repeated injury by needle injection is sufficient to induce a fibrotic phenotype in *Lair1^-/-^* mice. This was accompanied by increased skin collagen content in *Lair1^-/-^* mice in both NaCl (p<0.05) and bleomcyin (p=0.065) treatment groups (Figure 5E).

In the fibrotic phase of this model, (4 weeks), *Lair1^-/-^*mice continued to show increased skin thickness measured by caliper (Figure 5F) and collagen content (Figure 5G) compared to WT mice after bleomycin treatment, but not after NaCl injections. We further confirmed the increased dermal thickness in *Lair1^-/-^* mice challenged with bleomycin by performing histology with Masson’s trichrome staining (Figure 5H). By immunohistochemistry (IHC), we found an accompanying increase in αSMA+ dermal myofibroblasts (Figure 5I).

The increased fibrotic response in *Lair1^-/-^* mice supports a non-redundant role of LAIR-1 in tissue repair and regulation of fibrosis. Absence of proper LAIR-1 ligation due to collagen degradation in fibrotic tissue could thus lead to continued propagation of the tissue repair response in diseases such as SSc.

## Discussion

In this study, we show that LAIR-1 represents an essential control mechanism for tissue repair. LAIR-1 function can be obstructed by the production of decoy ligands in fibrotic tissue, potentially perpetuating the fibrotic process in diseases like SSc.

LAIR-1 expression and intrinsic function was comparable in cells from SSc patients and HC, which is in line with our previous findings, that under inflammatory conditions LAIR-1 is expressed and upon ligation still efficiently reduces cytokine production^26^. In addition, the concentration of soluble LAIR-1 antagonists in the circulation of SSc patients is similar to HC, in contrast to the increased concentration of sLAIR-1 and LAIR-2 observed in rheumatoid arthritis patients and in patients with autoimmune thyroid diseases^34,35^.

Despite normal LAIR-1 expression and function in cells from SSc patients, an inadequate ligand might undermine its inhibitory function. Collagen synthesis is highly upregulated in SSc tissues, leading to compensatory collagen degradation mechanisms. Several MMPs, including MMP7, MMP9 and MMP12, are increased in SSc patients^36,37^, translating to increased concentrations of collagen degradation products in the circulation and in the skin, suggesting a high collagen turnover in this disease^38–40^. However, despite increased collagen degradation, excessive collagen production still results in an imbalance and enormous accumulation of collagen in tissues and organs of these patients^36,39–41^.

We now report that even though LAIR-1 can bind to the fibrotic skin of SSc patients and to the ECM deposited by fibroblasts under pro-fibrotic conditions, it cannot be functionally activated by the collagen products that are formed. Thus, these structures may act as decoy ligands and compete with intact collagen to prevent inhibitory signaling by LAIR-1. In contrast, larger collagen fragments, ranging from 30-75 kDa, have the ability to prevent the formation of osteoclasts and to inhibit T cell response through a LAIR-1-dependent mechanism^42,43^.

Conversely, smaller fragments do not exhibit this inhibitory effect on osteoclast formation^42^. This suggests that the length of the fragment plays a critical role in activating LAIR-1, and that potentially small collagen fragments could act as decoy molecules. Interestingly, gelatin, which is a mixture of peptides and proteins produced by the partial hydrolysis of collagen consists of small collagen fragments and also has the capacity to bind LAIR-1 but not to activate it. We previously found that LAIR-1 binding to trimeric collagen peptides is influenced by glycine-proline-hydroxyproline (GPO) content of collagen, and recently it was shown that citrullinated collagen binds LAIR-1, but does not signal, resulting in inactive LAIR-1^44^, although other factors contribute to the interaction as well^18,29^. In addition, we identified a collagen peptide (II-17) that binds to LAIR-1 but is unable to induce LAIR-1 signaling^29^. Thus, biochemical and structural alterations of collagen determine the LAIR-1 functional response.

The inadequate activation of LAIR-1 by fibrotic collagen could be partly explained by the lack receptor clustering in analogy to glycoprotein VI (GPVI). GPVI is a collagen activating receptor on platelets and the collagen-binding domain is homologous of LAIR-1^29,45–47^. The proximity of GPVI dimer-binding sites on the collagen fiber surface enables clustering of GPVI dimers, increasing avidity and bringing together the necessary components to initiate receptor signaling.^48,49^ LAIR-1 may have similar requirements for signaling. While short collagen degradation products may be sufficient for LAIR-1 binding, signaling may require longer collagen fibers containing a greater number of binding sites for LAIR-1 clustering inducing a downstream signaling cascade.

The production of the fibrotic collagen products in our *in vitro* system upon repeated exposure to TGFβ and CXCL4 is dependent on PDGF receptor signaling and MMP activity. PDGF family members are potent mitogenic factors and are involved in chemotaxis and cell migration^32^. Both PDGFs and PDGFRs are upregulated in skin lesions of SSc patients and this pathway promotes myofibroblast activation^50^. In addition, PDGF proteins, particularly PDGF-D, upregulate the expression of multiple MMPs^51,52^, and are consequently implicated in collagen degradation^53^. Collagen fragments, particularly 3/4 length collagen fragments, in turn enhance PDGF migratory properties^54^, which may contribute to the increased motility we observed upon TGFβ plus CXCL4 stimulation. Indeed, MMP inhibition reduced cell motility, in agreement with previous work^55^.

The presence of collagen degradation products in SSc, as result of MMP activity and PDGF signaling as our data indicate, may lead to an inefficient LAIR-1 inhibitory signal in immune cells and thus perpetuate inflammation and consequent fibrosis in these patients.

In mice, the absence of LAIR-1 resulted in an increased presence of αSMA^+^ myofibroblast and increased collagen deposition and dermal thickness, not only in response to the fibrosis-promoting agent bleomycin, but also in response to the repetitive skin injury caused by the control injections. Thus, LAIR-1 deficient mice are more sensitive to developing fibrosis upon injury. This finding was consistent throughout experiments, although we did observe variability in the extension of the increased skin fibrosis in LAIR-1 deficient mice, reflected either by skin caliper, collagen content or histology, which may have been due to differences in the efficacy of bleomycin inducing skin fibrosis as well as different operators performing the model in different experiments^56^.

Since LAIR-1 can regulate a large variety of immune cells, it is possible that the exacerbated fibrotic phenotypic observed in LAIR-1 deficient mice is driven by multiple immune cells, including monocytes, macrophages, DCs, T cells and B cells. In particular, inflammatory monocytes and tissue-resident macrophages highly express LAIR-1 and are key regulators that maintain tissue homeostasis and are important for both driving and resolving tissue repair^2,26,27^. Further studies are needed to determine which cell types are essential for LAIR-1 mediated regulation of fibrosis.

Taken together, we identified a crucial role for LAIR-1 in the regulation of tissue repair using LAIR-1 deficient mice and show that in response to soluble factors that drive inflammation and fibrosis in SSc, human fibroblasts generate large amounts of degraded collagen, with impaired capacity to induce LAIR-1 signaling. Our data may explain how dysregulation can occur in fibrosis resulting in a vicious cycle: LAIR-1 normally senses collagen thus providing negative feedback to further collagen production, while collagen fragments, presumably generated during injury, prevent the negative feedback so that new collagen can be made to replace degraded collagen. However, collagen fragments produced under inflammatory conditions can remove this negative feedback even when there is excess collagen present in fibrotic tissues (Supplemental Figure 6). Thus, a circuit designed to maintain homeostasis has a vulnerability to dysregulation resulting in fibrosis.

Since LAIR-1 is expressed and intrinsically functional in SSc patients, our study opens novel avenues for the treatment of SSc and perhaps other fibrotic diseases by harnessing the LAIR-1 feedback loop to halt the cascade of inflammation and fibrosis.

## Materials and methods

### Patients

Blood from SSc patients was obtained from the University Medical Center Utrecht and Maasstad Hospital Rotterdam. Early SSc (eaSSc) patients were classified according to Leroy and Medsger^57^. Definite SSc patients fulfilling the classification the American College of Rheumatology/European League Against Rheumatism 2013 classification criteria^58^ without manifesting skin fibrosis were defined as non-cutaneous SSc (ncSSc); fibrotic SSc patients were stratified to limited cutaneous (lcSSc) or diffuse cutaneous SSc (dcSSc) according to the extent of skin involvement^59^ (Supplementary table 1). Blood from sex- and age-matched healthy controls (HC) was obtained from the Mini-Donor Service of the University Medical Center Utrecht. The study was conducted according to the Helsinki declaration and it was approved by the local institutional medical ethics review board. All subjects provided written informed consent. Samples and clinical information were treated anonymously immediately after collection.

### Animal Experiments

*Lair1*^−/−^ mice were generated on the C57BL/6 background by Taconic Artemis as described by Smith *et al* .^60^ Female *Lair1^-/-^* and wild-type (WT) mice derived from the same breeding line and housed under the same conditions were used for this study. Animal experiments were approved by the Committee on Animal Experiments of the Utrecht University and performed at the Central Animal Laboratory, Utrecht University.

### Bleomycin mouse model of SSc

WT and *Lair1*^−/−^ female mice were treated daily with subcutaneous injections into the back during either 2 weeks (1 independent experiment, n=6 per group, age ±28 weeks,) or 4 weeks (1 independent experiment, n=6 per group, age ±12 weeks; 3 independent experiments, n=3-6 per group, age ±28 weeks) of bleomycin (200 µL, 1000 µg/mL or 500 µg/mL respectively, Bleomedac - Medac or Baxter). Two days after the last injections, mice were sacrificed by cervical dislocation.

### Histochemical Analysis

Skin sections (5 μm) of paraffin-embedded tissue were stained with Mason’s trichrome, to evaluate collagen content and organization. For αSMA staining, deparaffinized sections were rehydrated, and streptavidin/biotin blocking kit (Vector Laboratories) was applied, followed by blocking step with normal goat serum (Cell Signaling Technology). Next, samples were stained with rabbit-anti mouse αSMA (2 µg/ml, Abcam); followed by a goat-anti rabbit antibody-biotin (1:300, Vector Laboratories) and subsequent Vectastain ABC HRP kit (Vector Laboratories). ImmPact NovaRED (Vector Laboratories) was used as substrate. Samples were counterstained with Mayer’s hematoxylin. Images were acquired on an Aperio Digital Pathology Slide Scanner (Leica Biosytems). Images were analysed using the Aperio ImageScope software (Leica Biosytems) and dermal thickness was measured according to the protocol described by Błyszczuk *et al*.^56^

### Quantification of Tissue Collagen

Collagen content was quantified by colorimetric assays from 4 mm skin punch biopsies. Skin sections were hydrolyzed and collagen content was determined in supernatants by QuickZyme total collagen assay (QuickZyme Bioscience) or the Total Collagen Assay Kit (Abcam), according to the manufacturer’s instructions.

### Western blotting

Murine skin samples were harvested from the injections areas and immediately stored in RNA*later* stabilization solution (Invitrogen – Thermo Fisher Scientific) until further analysis. Protein was extracted using the NucleoSpin RNA/Protein kit (Macherey – Nagel) according to the manufacturer’s instructions. After protein precipitation, washing and drying steps, protein samples were denatured in PSB-TCEP (Protein Solving Buffer containing reducing agent). Equivalent amounts of total protein lysate were subjected to electrophoresis on NuPAG 4–12% Bis-Tris Protein Gels (Thermo Fisher Scientific) and proteins were transferred to polyvinylidene difluoride membranes (Millipore). Membranes were incubated overnight at 4°C with the specific primary antibodies anti-αSMA and anti-tubulin (Supplementary table 2). Membranes were then washed and incubated in tris-buffered saline / Tween 20 containing horseradish peroxidase–conjugated secondary antibody. Protein was detected with SuperSignal^tm^ West Femto maximum sensitivity substrate (Thermo Scientific) using a ChemiDoc MP System (Biorad). Densitometry analysis was performed with Image J software. Relative αSMA protein expression was normalized to tubulin expression.

### Cell isolation

Peripheral blood was obtained from HC and SSc patients in lithium heparin tubes. Peripheral blood mononuclear cells (PBMCs) were isolated by density centrifugation using Ficoll-Paque Plus (GE Healthcare). Plasmacytoid dendritic cells and monocytes were isolated using anti-BDCA4 or anti-CD14 magnetic microbeads (Miltenyi Biotec) respectively, based on positive separation on auto-MACS assisted cell sorting (Miltenyi Biotec) according to the manufacturer’s protocol.

### Flow cytometry analysis

LAIR-1 expression on PBMCs was determined on freshly isolated cells, blocked with normal mouse serum (Fitzgerald) and then stained for 20 min at 4°C with fluorochrome-conjugated monoclonal antibodies according to the panel on Supplementary table 2. Samples were acquired on a BD LSR Fortessa (BD Biosciences) using the BD FACSDiva software (BD Biosciences). FlowJo software (Tree Star) was used for data analyses (analysis strategy in Supplemental Figure 7A-B).

### Immunohistochemistry staining for LAIR-1

Skin frozen sections (6 μm) from HC and SSc patients samples were fixed in 10% neutral buffered formalin for 15 sec. at room temperature (RT). After washing step with PBS+0.05% Tween-20, 0.3% Hydrogen peroxide was applied to the specimens, that were then blocked with 2.5% normal horse serum (Vector Laboratories) diluted in PBS. Next, 10 µg/mL of either mouse anti-human LAIR-1 (clone Dx26; purified in house) or mouse isotype control IgG1 (Dako) diluted in PBS+1% BSA buffer were incubated overnight at 4°C. Samples were then washed and incubated for 30 min with anti –mouse Ig HPR (Vector Laboratories). After washing, AEC+ Substrate-Chromogen (Dako) was used as substrate. Samples were counterstained with Mayer’s hematoxylin. Image acquisition was performed on an Olympus BX41 microscope. Images were further processed with ImageJ software. A representative LAIR-1 and isotype control staining is shown in Supplemental Figure 8A.

### Analysis of LAIR-1 function

LAIR-1 inhibitory function was assessed on purified pDCs and purified monocytes from HC and SSc patients as described before.^26^ Briefly, 96 well Nunc culture or 24 well Nunc culture plates (Thermo Fisher Scientific) were coated with 10 µg/mL of anti-LAIR-1 agonist mAb (clone Dx26) or 10 µg/mL of mouse isotype control IgG1 (eBioscience-Thermo Fisher Scientific) diluted in PBS overnight at 4°C. A total of 5.0×10^4^ pDCs or 5.0×10^5^ monocytes were seeded in the pre-coated plates with RPMI 1640 GlutaMAX (Life Technologies-Thermo Fisher Scientific), supplemented with 10% heat-inactivated fetal calf serum (FCS, Biowest Riverside) and 1% penicillin/streptomycin (Thermo Fisher Scientific), after incubation with Fc receptor blocking reagent (Miltenyi Biotech), for 2 hours at 37°C in a 5% CO_2_ incubator. Then, pDCs were either left unstimulated or stimulated with CPG-C-TLR9 ligand (500 µM) and monocytes were either left unstimulated or stimulated with LPS-TLR4 ligand (100 ng/mL), all from Invivogen. Cells were incubated overnight at 37°C in a 5% CO_2_ incubator and afterwards supernatants were harvested and stored at -80°C until further analysis.

### Measurement of cytokine production

Cytokines in cell-free supernatants were measured using enzyme-linked immunosorbent assay (ELISA) for IL-6 (Sanquin), TNF-α (Diaclone) and IFN-α (eBiosience, Thermo Fisher Scientific), following the manufacturer’s instructions. The cytokine production in monocytes and pDCs from HC and SSc patients upon stimulation is shown in Supplementary table 3. Soluble (s)LAIR-1 (as described previously^34^) and LAIR-2 (LifeSpan BioSciences) were also measured by ELISA in the serum of HC and SSc patients.

### LAIR-1 - Collagen binding assay

Collagen binding to LAIR-1 stably transduced K562-cells^18^ was assessed by flow cytometry adapted from the previously described protocol.^61^ In brief, collagen I FITC labelled was incubated for 1 hour at room temperature with serum either from SSc patients or HC (1:30 dilution). Next, 1.5×10^5^ K562-cells overexpressing LAIR-1 were incubated with the collagen-serum mix for 30 min. After a washing step, cells were incubated with anti-LAIR-1 PE (clone Dx26, BD) for 15 min and then washed. Samples were acquired on a BD LSR Fortessa (BD Biosciences) using the BD FACSDiva software (BD Biosciences). The percentage (%) of cells binding to collagen was determined using the FlowJo software (Tree Star).

### Human skin staining with hLAIR-1-IgG

Frozen tissue sections (5 µm) from SSc and HC skin that were fixed in 10% acetone in the presence of 1.5% H_2_O_2_. Free biotin in the sections was blocked using the streptavidin/biotin blocking kit (Vector Laboratories). Tissue sections were stained with biotinylated hLAIR-1–IgG (a chimeric protein obtained by fusing the extracellular domain of human(h)LAIR-1 to the Fc region of human IgG1^62^), in the presence of 2% FCS. After washing with PBS, sections were treated with streptABComplex/HRP (Vector Laboratories) according to the manufacturer’s instructions, followed by AEC+ substrate-chromogen (Dako) and counterstained with hematoxylin. Images were acquired on an Olympus BX41 microscope and to quantify the hLAIR1-IgG staining, HALO (Indica Labs) Area Quantification-v1 algorithm was used with thresholds for positive and negative signals set based on controls. A classifier was built to identify the dermis and used within the analysis algorithm to quantify IHC signal and reduce the background from pigmented skin samples.

### Dermal fibroblasts culture

Dermal fibroblasts were isolated from 3–4-mm skin biopsy sections obtained from a clinically affected area of SSc patients, while HC dermal fibroblasts were obtained from skin biopsy sections as resected material after cosmetic surgery. Dermal fibroblast isolation was performed using a whole skin dissociation kit (Miltenyi Biotec) following the manufacturer’s instructions, and fibroblasts were routinely maintained in Dulbecco’s modified Eagle’s medium (DMEM; Invitrogen) supplemented with 10% FBS and 1% penicillin–streptomycin. Cells were used for experiments between passages 3 and 5.

### *In vitro* fibrotic model

Nunc Lab-Tek II chamber slides (ThermoFisher Scientific) or µ-Slide 8 Well Glass Bottom (IBIDI) were precoated with 0.001% poly-l-lysine (Sigma-Aldrich), washed with PBS and air-dried. Dermal fibroblasts were seeded in DMEM containing 10% FBS for 24 hours and then incubated overnight in DMEM containing 0.1% FBS. Cells were then stimulated with recombinant TGFβ2 (10 ng/mL, R&D Systems), recombinant CXCL4 (5 µg/mL, Peprotech) or the combination of both in DMEM containing 2% FBS and 50 μg/ml of L-ascorbic acid (Wako Pure Chemical), and refreshed 2 and 4 days later. Where indicated, PDGFR inhibitor CP-868596/Crenolanib (Selleckchem) and the metalloproteinase inhibitor GM6001/ Galardin (Merck Millipore) were used at the concentration of 1 μM and 100 µM respectively and refreshed as described above. In some experiments cells under stimulation were imaged live on the IncuCyte system (Essen BioScience). After 6 days of stimulation, or otherwise indicated, cells were used for immunofluorescence, electron microscopy analysis or reporter cell assay.

### Immunofluorescence

Cells were fixed with 4% paraformaldehyde, washed with PBS–1% bovine serum albumin (BSA), and blocked in PBS containing 5% normal serum and 1% BSA. Cells were incubated with biotinylated hLAIR-1-IgG, anti-collagen I, or pan-anti-collagen antibody mix (anti-collagen I, II, III, IV, V) for 1 hour at room temperature, washed, and incubated with secondary antibodies conjugated with Alexa Fluor 488, Alexa Fluor 594 or Alexa Fluor 647 for 30 minutes at room temperature (Supplementary table 2 for antibody information).

The collagen hybridizing peptide (CHP, 3Helix) staining was performed according to the manufacturer’s protocol. Briefly, cells were fixed with 4% paraformaldehyde and washed with PBS+1% BSA. Next, samples were blocked with 5% normal goat serum and 1% BSA and then incubated for 2 hours at 4°C with biotinylated CHP (2.5 µM), previously denatured at 80°C for 5 min (non-denatured CHP was used to assess unspecific binding) plus LAIR-1-IgG (5 µg/mL). After washing with PBS+1% BSA, samples were incubated 1 hour at room temperature with unlabeled anti-LAIR-1 mAb (2.5 µg/mL, clone Dx26) followed by a last step incubation with streptavidin AF647 and goat anti-mouse AF488 (both 2 µg/mL).

Cells were incubated with DAPI (Sigma-Aldrich) for 15 min at room temperature and finally washed with PBS.

Imaging data were acquired on a Zen2009 LSM 710 confocal microscope (Zeiss) or on the IncuCyte system (Essen BioScience). For the quantification of LAIR-1-IgG and Collagen I co-staining the .lsm files were converted to .czi files and then analyzed using HALO (Indica Labs) Area Quantification FL v1 algorithm. Positive and negative samples were used to set thresholds for each marker. To quantify the areas of plaques (hLAIR-1-IgG binding structures), HALO classifiers were trained on negative and positive samples. Separate classifiers were developed for images captured on the Incucyte versus the Zen2009 LSM 710 confocal microscope.

### Electron microscopy

Dermal fibroblasts were grown on carbon coated glass coverslips as described above, fixed with 4% Formaldehyde, labelled with pan-collagen mix of antibodies antibody, hLAIR-1-IgG fusion protein, and counterstained with DAPI as also stated above. The coverslips were imaged in the fluorescence microscope (Thunder, Leica) to confirm the presence of LAIR-1 binding ECM structures. After fluorescence microscopy, cells on coverslips were fixed in half strength Karnovsky fixative (2.5% Glutaraldehyde (EMS) + 2% Formaldehyde (Sigma)) pH 7.4 at RT for 2 h. Cells were rinsed and stored in 1M Phoshate Buffer pH 7.4 at 4°C until further processing. Post-fixation was performed with 1% OsO4, 1.5% K3Fe(III)(CN)6 in 1M Phoshate Buffer pH 7.4 for 2 h. Cells were then dehydrated in a series of aceton, and flat embedded in Epon (SERVA) similar to reported before.^63^ After curing, the coverslips were detached from the resin block and ultrathin sections were cut (Leica Ultracut UCT), collected on formvar and carbon coated transmission electron microscopy (TEM) grids, and stained with uranyl acetate and lead citrate (Leica AC20). Micrographs were collected on a JEM1010 (JEOL) equipped with a Veleta 2k×2k CCD camera (EMSIS, Munster, Germany) or on a Tecnai12 (FEI Thermo Fisher) equipped with a Veleta 2k×2k CCD camera (EMSIS, Munster, Germany) and operating SerialEM software.

### Binding assay

96-well black flat-bottom plates (Corning) were coated overnight at 4 °C with purified collagen I from human placenta (Sigma), gelatin from bovine skin (Sigma) or BSA (for unspecific binding) diluted in PBS (50 µL/well). After washing with PBS, proteins were fixed with 100% ice-cold methanol at -20°C for 30 min and then washed. Wells were then blocked for 30 min with PBS containing 1% of normal donkey serum and 2% FBS. Blocking solution was discarded and replaced with hLAIR-1-IgG biotinylated (5 µg/mL) and incubated for 1 hour at RT. After washing, wells were incubated with a Streptavidin AF647 (5 µg/mL) for 1 hour at RT. After washing, imaging and data acquisition was performed on a LI-COR - Odyssey system. Signal intensity was obtained for each well.

### Reporter cell assay

2B4 T cell hybridoma cells with stable expression of a NFAT-GFP reporter and hLAIR-1-CD3ζ or WT-CD3ζ were cultured as previously described.^18^ To assess hLAIR-1-CD3ζ response to either purified collagen I or gelatin, 96-well black flat-bottom plates (Corning) were coated overnight at 4 °C with purified collagen I from human placenta (Sigma), gelatin from bovine skin (Sigma) or BSA (for unspecific binding) diluted in PBS (50 µL/well). After washing, 5.0× 10^4^ cells in 200 μL were added to each well, and plates were incubated at 37°C for 20 h and analyzed for GFP expression by flow cytometry.

To evaluate hLAIR-1-CD3ζ response to fibrotic ECM, dermal fibroblasts cultured and stimulated as described above, were rinsed with PBS and 1.5×10^5^ hLAIR-1-CD3ζ - NFAT-GFP reporter cells were added. For the experiment controls, µ-Slide 8 Well Glass Bottom (IBIDI) were coated overnight at 4°C with purified collagen I, BSA, or anti–hLAIR-1 mAb (clone Dx26) at 5 μg/mL in PBS. After washing, cells were added to each well. Cells were incubated for 20 hours and GFP expression was measured by flow cytometry. In parallel, samples were incubated in the IncuCyte system and live imaging was performed for 70 hours to determine GFP expression. Efficiency of the LAIR-1-CD3ζ and WT-CD3ζ response are shown in Supplemental Figure 9.

### LAIR-1 expression from profiling data

LAIR-1 gene expression was retrieved from array profiling data available on the Gene Expression Omnibus (GEO – NCBI) using GEO2R (NCBI).

### Statistics

Statistical analysis was performed using GraphPad Prism 8 software (GraphPad Software Inc.). Differences between experimental groups were analyzed using parametric paired t-tests, one-way ANOVA test or non-parametric Mann–Whitney U test or Wilcoxon’s test, when appropriate. Pearson’s correlation coefficient test was applied to detect the association between different parameters. Two-sided testing was performed for all analyses. Differences were considered to be statistically significant at p<0.05

## Acknowledgements

We thank to Department of Rheumatology & Clinical Immunology in University Medical Centre Utrecht and to the Department of Rheumatology and Clinical Immunology, Maasstad Hospital Rotterdam for patient inclusion. We would also like to thank Karin de Cortie (University Medical Center Utrecht) for the technical support with the animal experiments.

## Competing interests

DF, MR and MEMG were full time employees of Boehringer Ingelheim. TRDJR was a principal investigator in the immune catalyst program of GlaxoSmith-Kline, which was an independent research program. He did not receive any financial support. Currently, TRDJR is an employee of Abbvie where he holds stock. TRDJR had no part in the design and interpretation of the study results after he started at Abbvie. LM’s research lab has received financial support for investigator-initiated studies from Boehringer Ingelheim, NextCure and NGM biopharmaceuticals. The remaining authors declare that the research was conducted in the absence of any commercial or financial relationships that could be construed as a potential conflict of interest.

## Funding

TC was supported by a grant from the Portuguese national funding agency for science, research, and technology: Fundação para a Ciência e a Tecnologia [SFRH/BD/93526/2013]. S.G. is supported by the Miguel Servet program (CP19/00005) from the Instituto de Salud Carlos III (ISCIII) and the European Social Fund (“Investing in your future”). LM is supported by the Dutch Research Council (NWO) (Vici 918.15.608). The electron microscopy within this work is part of the research program National Roadmap for Large-Scale Research Infrastructure (NEMI), financed by NWO (184.034.014) to JK. NL is financed by ZonMw TOP grant (40-00812-98-16006) to JK and Anna Akhmanova. Part of this study was supported by an investigator-initiated collaborative research agreement between LM’s research lab and Boehringer Ingelheim.

## Author contributions

TC, WM, TRDJR and LM were involved in study conception. TC, WM, SG, MIPR, JK, NL, TRDJR and LM designed the experimental approach and interpreted the data. TC, WM, SG, MIPR, AJA, ASM, BG, RGT, EE, MLM and TV carried out the experiments. TC, DF, TV and LV performed data analysis. AO and MC collected and supplied clinical data. MR and MEMG reviewed data. TC and LM wrote the original draft. All authors critically revised the manuscript and approved the final version.

## Ethics

Blood samples from SSc patients were obtained from the University Medical Center Utrecht and Maasstad Hospital Rotterdam. Skin samples were obtained from patients with SSc from the University Medical Center Utrecht. Blood from healthy controls was obtained from the Mini Donor Service of the University Medical Center Utrecht. The study was conducted according to the Helsinki declaration and ethical approval was requested and obtained from the Medical Ethical Research Committee in University Medical Center Utrecht and Maasstad Hospital Rotterdam. All subjects provided written informed consent. Samples and clinical information were treated anonymously immediately after collection.

Animal experiments were approved by the Committee on Animal Experiments of the Utrecht University and performed at the Central Animal Laboratory, Utrecht University.

## Notes

### Summary of Updates

Results section and Figure 5 were updated. Discussion section updated to clarify.

