## Supplemental materials for "Impaired LAIR-1-mediated immune control due to collagen degradation in fibrosis"

### Supplemental data

#### Supplemental Figure 1

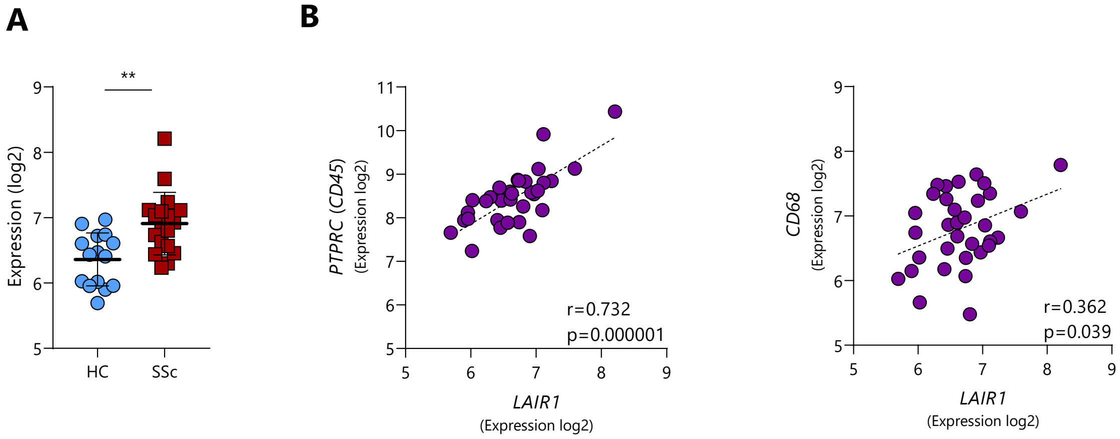


**Supplemental Figure 1. *LAIR1* expression in affected skin from SSc patients. (A)** *LAIR1* gene expression in skin from HC and SSc patients by array profiling and retrieved from publicly available dataset GSE95065. Results are represented as mean with SD. Differences were considered statistically significant between HC and SSc patients when $^{**}p < 0.01$ (Unpaired t-test) **(B)** Correlation between *LAIR1* expression and *PTPRC* (gene for CD45) or *CD68* expression in skin from HC and SSc patients (GSE95065). Correlations were assessed by the Pearson's correlation coefficient test.

#### Supplemental Figure 2

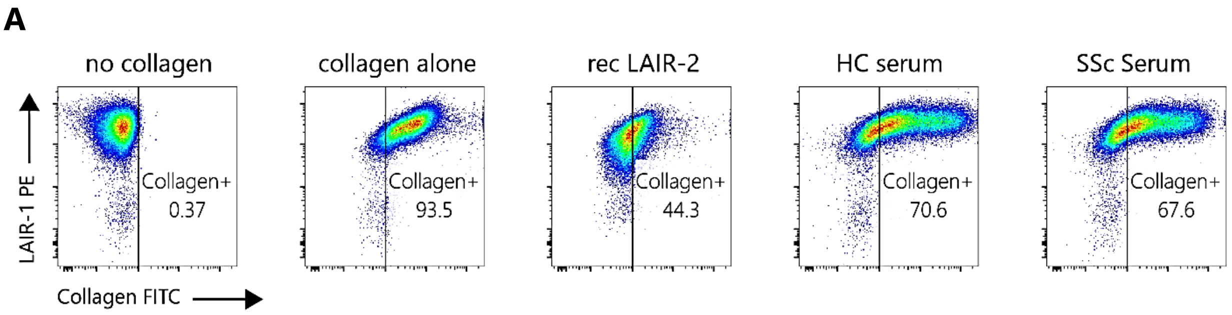


**Supplemental Figure 2. Collagen – LAIR-1 blocking with HC and SSc patients sera.**

**(A)** Representative flow cytometry analysis showing K562 overexpressing- LAIR-1 cells incubated either with FITC labelled collagen alone, or pre-incubated with recombinant LAIR-2 (40 µg/mL), or serum from HC or SSc patients.

**Supplemental Figure 3**

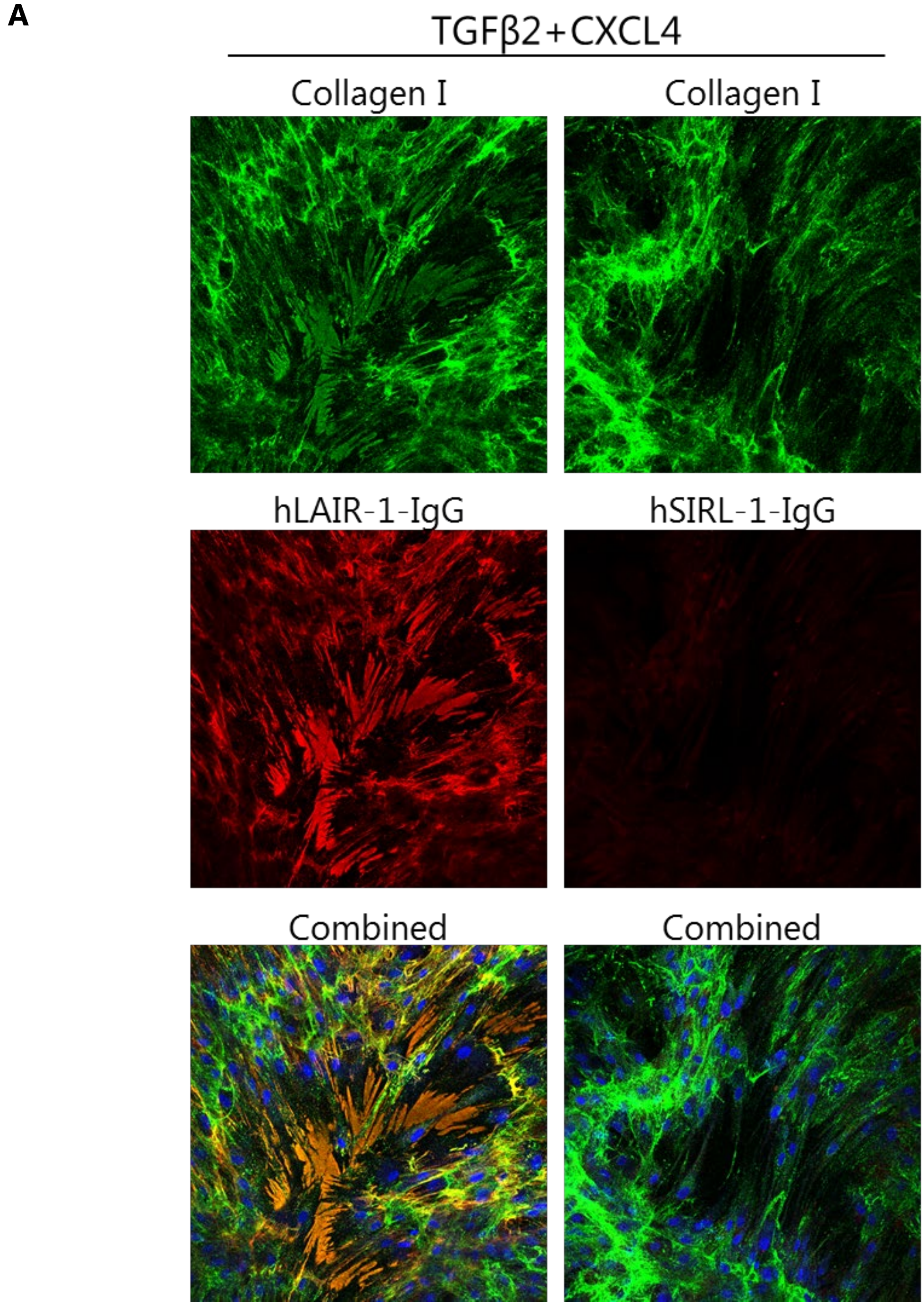


**Supplemental Figure 3. Immunofluorescence staining of ECM with hLAIR-1-IgG or hSIRL-1-IgG. (A)** ECM produced by HC fibroblasts upon TGFβ2 plus CXCL4 combination, stained with the pan-collagen mix of antibodies (green) either with hLAIR-1-IgG (red) or human Signal inhibitory receptor on leukocytes-1 (hSIRL-1)-IgG (red) for irrelevant binding control.

#### Supplemental Figure 4

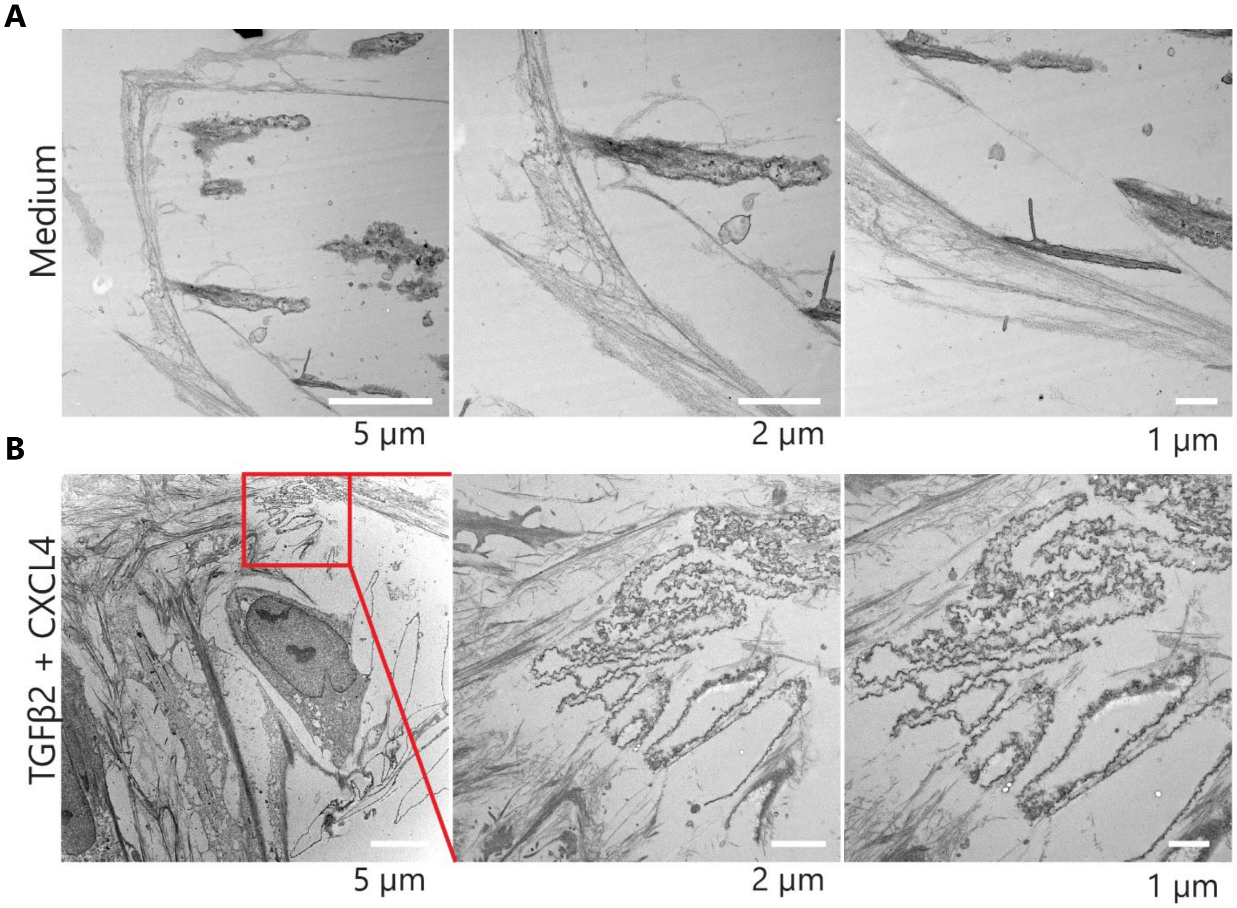


**Supplemental Figure 4. Transmission Electron Microscopy (TEM) images of ECM produced by fibroblasts stimulated with TGFβ2 in combination with CXCL4.** Additional TEM images of collagen fibers in **(A)** medium control (upper panel) and **(B)** TGFβ2 plus CXCL4 treated (lower panel) fibroblasts. Red square and corresponding zoom-in images show the ECM structure composed of agglomerated collagen fibers observed in TGFβ2 plus CXCL4 treated fibroblasts. Scale bars are 5 µm, 2 µm and 1 µm from left to the right in both conditions, respectively.

#### Supplemental Figure 5

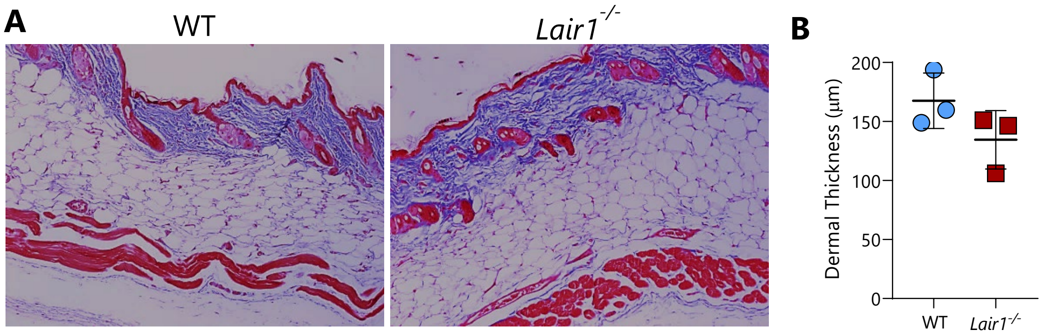


**Supplemental Figure 5. Histological analysis of skin from untreated WT and *Lair1$^{-/-}$* mice.** **(A)** Representative Masson's trichrome staining of skin from untreated WT and *Lair1$^{-/-}$* mice (C57BL/6J females, 66-64 weeks old) and **(B)** quantification of the dermal thickness (n=3 per group).

#### Supplemental Figure 6

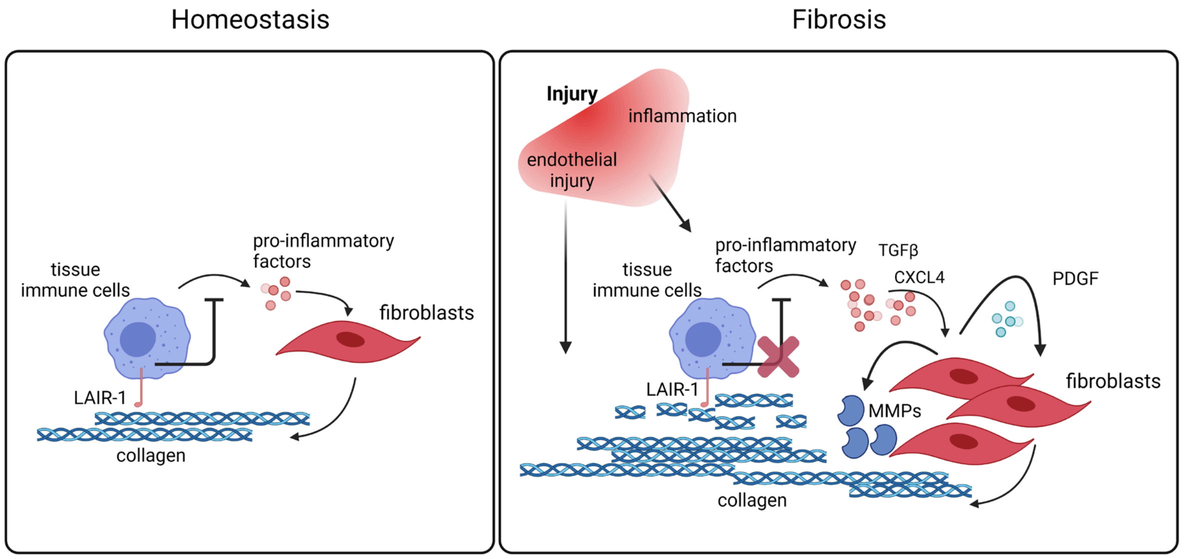


**Supplemental Figure 6. Proposed model for LAIR-1 role in fibrosis.** Tissue repair is a carefully regulated process, which entails a cascade of inflammation, repair and ultimately resolution events. Under homeostatic conditions, intact collagen acts as a ligand for LAIR-1, inhibiting tissue immune cells and leading to reduced production of inflammatory factors. In persistent inflammatory conditions, characterized by repetitive injury or endothelial damage, immune cells release substantial amounts of inflammatory mediators, which lead to fibroblasts hyperactivation, resulting in the excessive production of collagen. Simultaneously, there is an upregulation in the production of matrix metalloproteinases (MMPs), leading to the deposition of fragmented and disorganized collagen, that serves as a decoy ligand for LAIR-1, impeding its regulatory function and contributing to the persistence of inflammation and fibrosis.

#### Supplemental Figure 7

**A**

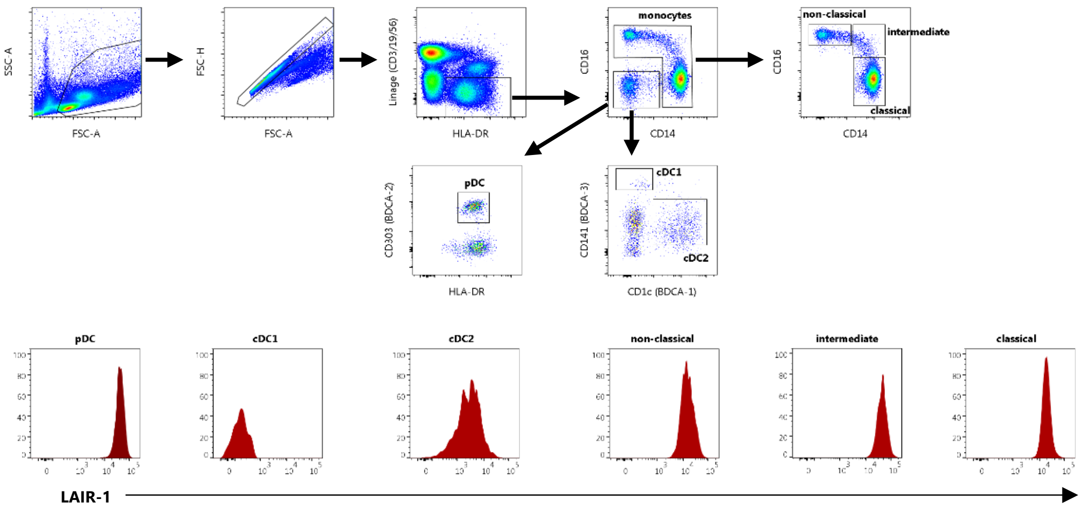


**B**

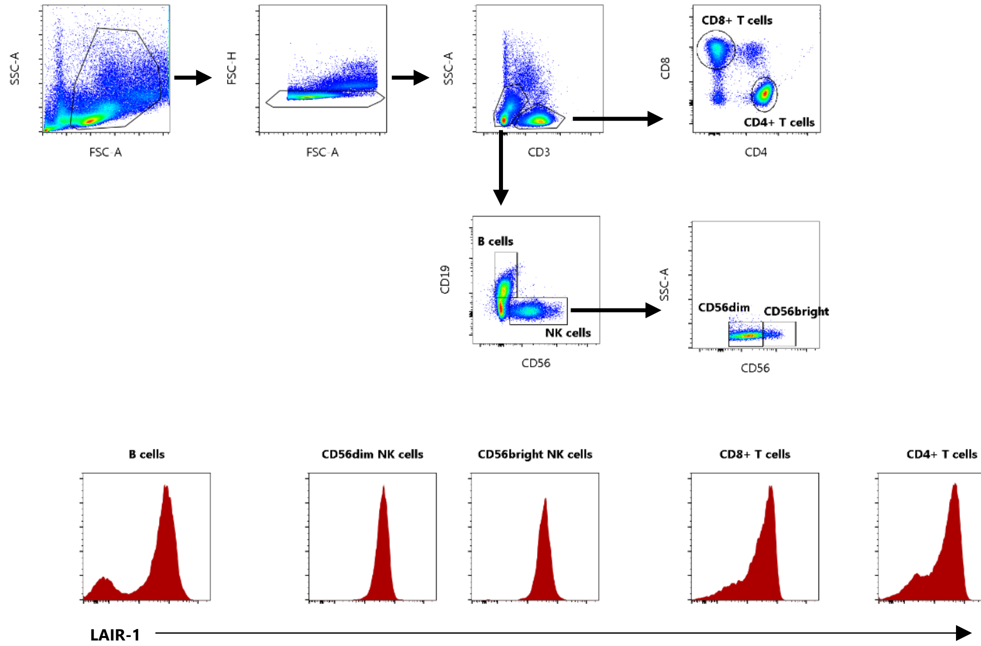


**Supplemental Figure 7. Flow cytometry analysis strategy to determine LAIR-1 expression on the different circulating immune cells of HC and SSc patients.** Flow cytometry gating strategy analysis to identify **(A)** plasmacytoid dendritic cells (pDC), conventional type 1 DCs (cDC1), conventional type 2 DCs (cDC2), non-classical, intermediate and classical monocytes, and **(B)** B cells,

$CD56^{bright}$ and $CD56^{dim}$ NK cells, CD8+ and CD4+ T cells, on peripheral blood mononuclear cells (PBMC) of HC and SSc patients.

#### Supplemental Figure 8

A

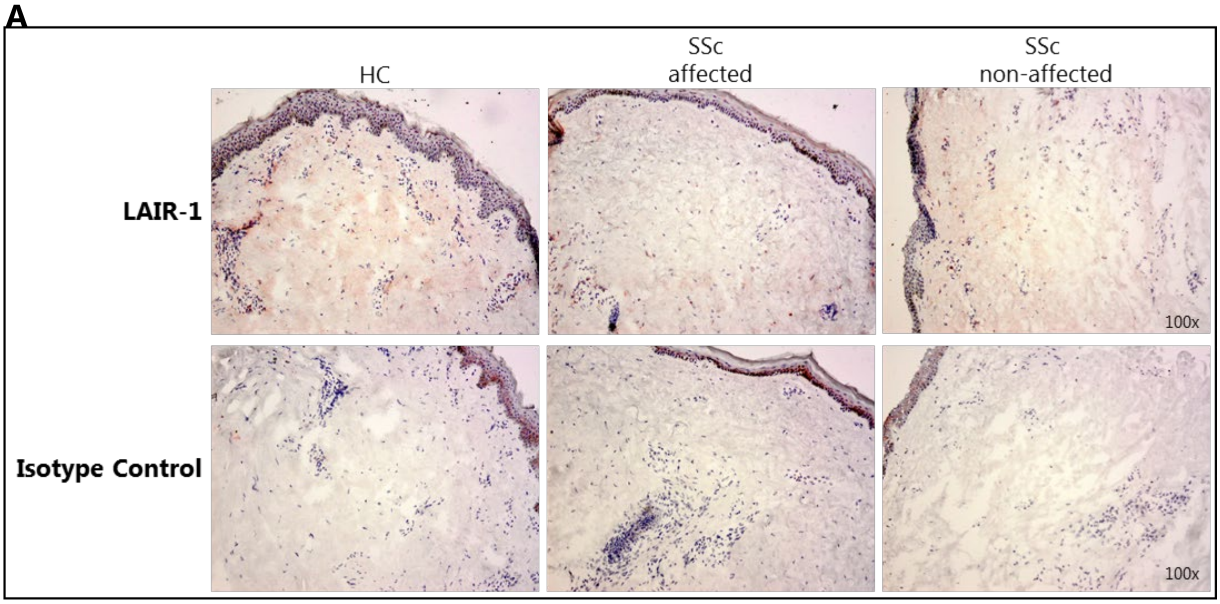


**Supplemental Figure 8. Immunohistochemistry analysis of LAIR-1 expression in HC and SSc skin. (A)** Representative images of LAIR-1 immunohistochemistry staining (red substrate) (top panel) and isotype control as negative control (bottom panel) in HC and SSc affected and non-affected skin (n=3 per group). Mayer's hematoxylin was used as nuclear counterstain.

#### Supplemental Figure 9

A

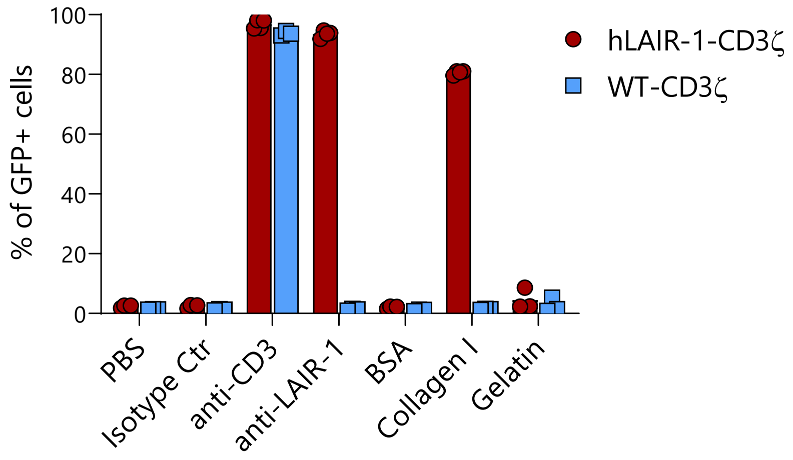


**Supplemental Figure 9. Efficiency of the LAIR-1 response using a NFAT-GFP reporter cell line transfected with hLAIR-1 and CD3ζ chimera.** **(A)** Response (% of GFP+ cells measured by flow cytometry) of hLAIR-1-CD3ζ or control WT-CD3ζ parental cells to PBS (background control), immobilized mouse IgG isotype control (5 µg/mL), anti-CD3 (5 µg/mL), anti-human LAIR-1 (Dx26) (5 µg/mL), BSA (5 µg/mL), purified human collagen I (5 µg/mL) or bovine skin gelatin (5 µg/mL) on a plastic culture plate.

**Supplemental Table 1.** SSc patients characteristics

| | n= | Age (years) | Female: *n* (%) | Early SSc | ncSSc | lcSSc | dcSSc |
|---|---|---|---|---|---|---|---|
| **LAIR-1 expression** (peripheral blood) | 26 | 53±14 | 17 (65%) | 4 (15.4%) | 6 (23.1%) | 13 (50%) | 3 (11.5%) |
| **LAIR-1 function** (peripheral blood) | 7 | 57±18 | 5 (71%) | 0 | 1 (14.3%) | 3 (42.9%) | 3 (42%) |
| **serum samples** (sLAIR-1 and LAIR-2 ELISA) | 30 | 54±18 | 21 (70%) | 8 (26.7%) | 3 (10%) | 11 (36%) | 8 (26.7%) |
| **serum samples** (K562-LAIR-1 – collagen blocking) | 36 | 55±10 | 22 (61%) | 2 (5.6%) | 6 (16.7%) | 19 (52.8%) | 9 (25%) |
| **skin samples** (IHC: LAIR-1 and hLAIR-1-IgG) | 9 | 43±8 | 7 (77%) | 0 | 0 | 4 (44.4%) | 5 (55.6%) |
| **dermal fibroblasts** (IF) | 6 | 48±14 | 4 (67%) | 0 | 0 | 2 (33.3%) | 4 (66.7%) |

**Supplemental Table 2.** Antibodies used in flow cytometry (FCM), immunohistochemistry (IHC) Immunofluorescence (IF) and functional assays.

| Target | Label | Manufacturer | Catalogue nr. | Dilution | Clone | Application |
|---|---|---|---|---|---|---|
| CD1c (BDCA1) | BV421 | Biolegend | 331526 | 1:20 | L161 | FCM |
| CD14 | APC-H7 | BD | 560180 | 1:50 | MφP9 | FCM |
| BDCA2 | PerCP-Cy5.5 | Biolegend | 354209 | 1:50 | 201A | FCM |
| BDCA3 | PECy7 | Biolegend | 344110 | 1:50 | M80 | FCM |
| CD3 | AF700 | Biolegend | 300424 | 1:25 | UCHT1 | FCM |
| CD19 | AF700 | eBioscience | 56-0199-42 | 1:50 | HIB19 | FCM |
| CD56 | AF700 | BD | 557919 | 1:50 | B159 | FCM |
| CD16 | V500 | BD | 561394 | 1:50 | 3G8 | FCM |
| HLA-DR | APC | BD | 559866 | 1:50 | G46-6 | FCM |
| CD3 | AF700 | Biolegend | 300424 | 1:40 | UCHT1 | FCM |
| CD4 | PB | Biolegend | 300521 | 1:50 | RPA-T4 | FCM |
| CD8 | V500 | BD | 560774 | 1:100 | RPA-T8 | FCM |
| CD45Ro | PECy7 | BD | 337168 | 1:20 | UCHL1 | FCM |
| CD27 | APC-eF780 | eBioscience | 47-0279-42 | 1:10 | O323 | FCM |
| CD19 | PerCP | Biolegend | 302228 | 1:100 | HIB19 | FCM |
| CD56 | APC | Biolegend | 318310 | 1:30 | HCD56 | FCM |
| LAIR-1 | PE | BD | 550811 | 1:20 | Dx26 | FCM |
| αSMA | Unlabelled | Abcam | ab5694 | 2 µg/mL/1:000 | Polyclonal | IHC/WB |
| Tubulin | Unlabelled | Sigma | T9026 | 1:2000 | DM1A | WB |
| Isotype control | Unlabelled | Abcam | ab27478 | 2 µg/mL | - | IHC |
| IgG (H+L) | Biotin | Vector Labs | BA1000 | 1:300 | Polyclonal | IHC |
| LAIR-1 | Unlabelled | Purified in house | - | 10 µg/mL | Dx26 | IHC/functional |
| mouse isotype | Unlabelled | Dako/Agilent | X0931 | 10 µg/mL | - | IHC |
| mouse isotype | Unlabelled | Invitrogen | 16-4714-85 | 10 µg/mL | | functional |
| Mouse CD3ε | Unlabelled | BD | 553057 | 5 µg/mL | 145-2C11 | functional |
| hLAIR-1-IgG | Biotin | Purified in house | - | 5-10 µg/mL | - | IHC/IF |
| hSIRL-1-IgG | Biotin | Purified in house | - | 5 µg/mL | - | IF |
| Collagen I | Unlabelled | SouthernBiotech | 1310-01 | 2 µg/mL | - | IF |
| Collagen-Pan | Unlabelled | Invitrogen | PA5-104252 | 5 µg/mL | Polyclonal | IF |
| Donkey anti-Goat IgG | Alexa Fluor 488 | Invitrogen | A11055 | 2 µg/mL | - | IF |
| Streptavidin | Alexa Fluor 594 | Invitrogen | S32356 | 2 µg/mL | - | IF |
| Streptavidin | Alexa Fluor 647 | Invitrogen | S32357 | 2 µg/mL | - | IF |
| Goat anti-Rabbit IgG (H+L) | Alexa Fluor 488 | Invitrogen | A11070 | 2 µg/mL | - | IF |
| Swine anti-Rabbit | HRP | DAKO | P021702 | 1:10.000 | - | WB |
| Goat anti-mouse | HRP | DAKO | P0447 | 1:10.000 | - | WB |

**Supplemental Table 3.** Levels of IL-6 and TNF-α produced by monocytes and IFN-α produced by plasmacytoid dendritic cells (pDCs).

Isolated cells were pre-treated with LAIR-1 agonist (Dx26) and monocytes were stimulated with LPS and pDC with CpC-C. Production of IL-6 and TNF-α by monocytes and IFN-α by pDCs was evaluated using ELISA. Data are shown as mean ± SD. Statistically significant differences between isotype control and anti-LAIR-1 conditions were considered when *p < 0.05 (Wilcoxon's test).

| | | | IL-6 (pg/mL) | TNF-α (pg/mL) | IFN-α (pg/mL) |
|---|---|---|---|---|---|
| **Monocytes** | | | | | |
| Medium | HC | Isotype Control | 9072±16252 | 1014±848.5 | - |
| | | anti-LAIR-1 | 1504±2622* | 1183±117 | - |
| | SSc | Isotype Control | 3950±6177 | 624.6±329.7 | - |
| | | anti-LAIR-1 | 4291±9306 | 710.9±676.4 | - |
| LPS TLR4- ligand | HC | Isotype Control | 76985±31227 | 19026±11134 | - |
| | | anti-LAIR-1 | 12177±9200* | 5678±1816* | - |
| | SSc | Isotype Control | 74469±47792 | 14714±8638 | - |
| | | anti-LAIR-1 | 24416±46928* | 9329±13105 | - |
| **pDCs** | | | | | |
| Medium | HC | Isotype Control | - | - | 479±237.8 |
| | | anti-LAIR-1 | - | - | 361.9±107.9 |
| | SSc | Isotype Control | - | - | 469.5±274 |
| | | anti-LAIR-1 | - | - | 327.9±111 |
| CpG-C TLR9- ligand | HC | Isotype Control | - | - | 24202±16481 |
| | | anti-LAIR-1 | - | - | 12606±10915* |
| | SSc | Isotype Control | - | - | 13714±7885 |
| | | anti-LAIR-1 | - | - | 5856±5864* |
