## Supplemental videos for "Impaired LAIR-1-mediated immune control due to collagen degradation in fibrosis": Supplemental video legends_Carvalheiro_et_al.pdf

### Supplemental videos legends:

**Supplemental video 1.** HC fibroblasts unstimulated (medium) were imaged with phase contrast every 6 hours during 6 days on an Incucyte live system (4x magnification).

**Supplemental video 2.** HC fibroblasts stimulated with TGF-β2 combined with CXCL4 were imaged with phase contrast every 6 hours during 6 days on an Incucyte live system (4x magnification).

**Supplemental video 3.** HC fibroblasts unstimulated (medium) were imaged with phase contrast every 6 hours during 6 days on an Incucyte live system (4x magnification).

**Supplemental video 4.** HC fibroblasts unstimulated (medium) in the presence of PDGF receptor inhibitor (Crenolanib) were imaged with phase contrast every 6 hours during 6 days on an Incucyte live system (4x magnification).

**Supplemental video 5.** HC fibroblasts stimulated with TGF-β2 combined with CXCL4 were imaged with phase contrast every 6 hours during 6 days on an Incucyte live system (4x magnification).

**Supplemental video 6.** HC fibroblasts stimulated with TGF-β2 combined with CXCL4 in the presence of PDGF receptor inhibitor (Crenolanib) were imaged with phase contrast every 6 hours during 6 days on an Incucyte live system (4x magnification).

**Supplemental video 7.** HC fibroblasts unstimulated (medium) in the presence of DMSO were imaged with phase contrast every 6 hours during 6 days on an Incucyte live system (4x magnification).

**Supplemental video 8.** HC fibroblasts unstimulated (medium) in the presence of MMPi inhibitor (GM6001) were imaged with phase contrast every 6 hours during 6 days on an Incucyte live system (4x magnification).

**Supplemental video 9.** HC fibroblasts stimulated with TGF-β2 combined with CXCL4 in the presence of DMSO were imaged with phase contrast every 6 hours during 6 days on an Incucyte live system (4x magnification).

**Supplemental video 10.** HC fibroblasts stimulated with TGF-β2 combined with CXCL4 in the presence of MMPi inhibitor (GM6001) were imaged with phase contrast every 6 hours during 6 days on an Incucyte live system (4x magnification).

**Supplemental video 11.** hLAIR1-CD3ζ reporter cells incubated on ECM produced by unstimulated HC fibroblasts. Images every 1 hour for 70 hours with phase contrast and fluorescence (GFP+ cells in green) on an Incucyte live system (10x magnification).

**Supplemental video 12.** hLAIR1-CD3ζ reporter cells incubated on ECM produced by unstimulated SSc fibroblasts. Images every 1 hour for 70 hours with phase contrast and fluorescence (GFP+ cells in green) on an Incucyte live system (10x magnification).

**Supplemental video 13.** hLAIR1-CD3ζ reporter cells incubated on ECM produced by HC fibroblasts stimulated with TGFβ2+CXCL4. Images every 1 hour for 70 hours with phase contrast and fluorescence (GFP+ cells in green) on an Incucyte live system (10x magnification).

**Supplemental video 14.** hLAIR1-CD3ζ reporter cells incubated on ECM produced by SSc fibroblasts stimulated with TGFβ2+CXCL4. Images every 1 hour for 70 hours with phase contrast and fluorescence (GFP+ cells in green) on an Incucyte live system (10x magnification).
